# Loss of HSC stemness identity is associated with exhaustion and hyporesponsiveness in GATA2 deficiency syndrome

**DOI:** 10.1101/2023.08.07.551811

**Authors:** Laetitia Largeaud, Vincent Fregona, Laura Jamrog, Camille Hamelle, Stéphanie Dufrechou, Naïs Prade, Esmaa Sellam, Pauline Enfedaque, Manon Bayet, Sylvie Hébrard, Mathieu Bouttier, Christine Didier, Eric Delabesse, Bastien Gerby, Marlène Pasquet, Cyril Broccardo

**Author notes:** contributed equally (Laetitia Largeaud, Vincent Fregona, Laura Jamrog).

## Abstract

Germline *GATA2* mutations lead to a syndrome involving both immunodeficiency and myeloid malignancies. Since GATA2 is a key player in hematopoietic initiation and development, we specify the impact of these germline mutations on hematopoietic homeostasis by generated a knock-in mouse model expressing the recurrent *Gata2* R396Q missense mutation. These mice exhibit a hematopoietic stem and progenitor cell (HSPC) compartment profoundly impacted with increased HSC number, decreased self-renewal potential and inability to respond to acute inflammatory stimuli. Moreover, mutated HSPCs are predisposed to be hyporesponsive, as evidenced by lower interferon signaling and enrichment of inflammatory stress signatures. Furthermore, a Gata2 allelic specific expression results in a molecular and functional heterogeneity of the mutated Long Term-HSC population. Altogether, we highlight that Gata2 plays a crucial role in the ability of HSCs to perceive and respond to their environment, and that germline mutation contributes to the decline in HSC functionality.

## Introduction

GATA2 is one of the master regulatory transcription factors of hematopoiesis, characterized by two zinc finger domains (N-ZF and C-ZF) which bind DNA and interact with protein partners to regulate its function^1^. During embryogenesis, GATA2 plays a crucial role in the generation and survival of the hematopoietic stem and progenitor cells (HSPCs)^2, 3^. Its deregulation leads to a decrease in the number of HSCs and a loss of their self-renewal capacity in young mice^4, 5^. Over the past decade, germline *GATA2* mutations have been extensively identified in patients with a wide range of clinical presentations such as hematological malignancies, immunological defects and vascular disorders^6–12^. The spectrum of germline *GATA2* mutations is broad in this syndrome^13^. Overall, frameshift, nonsense and enhancer +9.5 mutations suggest a haploinsufficiency mechanism of pathogenesis, while recurrent missense mutations may have aberrant activity^14, 15^. Moreover, patients with missense mutations are more prone to develop leukemic transformation, suggesting that these mutations may confer additional features^16^. However, their impacts remain poorly studied *in vivo* due to the very small number of available models^17, 18^. We therefore developed a *knock-in* mouse model of *Gata2* R396Q mutation and studied its impact on hematopoiesis.

Situated at the apex of hematopoietic hierarchy, long-term HSCs (LT-HSCs) have the ability to self-renew and differentiate to maintain hematopoietic homeostasis throughout the entire lifespan of an organism^19^. We report here that in contrast to the *Gata2*^+/-^ mouse model, *Gata2*^R396Q/+^ mice have an increased number of LT-HSCs from embryogenesis to old age. Despite this phenotype, challenging assays revealed functional defects explained at the molecular level by enrichment in the pathways of exhaustion and hyporesponsiveness. Moreover, it is now admitted that the HSC population is phenotypically and functionally heterogeneous^20^. Previously, Gata2 has been identified as a gene exhibiting allelic specific expression (ASE) in non-physiological contexts such as acute myeloid leukemia (AML)^21^ or GATA2 deficiency^22^. Here, we show that an aberrant population of LT-HSCs predominantly expresses the *Gata2*-mutated allele highlighting a link between the heterogeneity of the HSC population and the ASE of *Gata2* gene.

Thus, our data shed light on the impact of a *Gata2* missense mutation on the hematopoietic tissue, allowing us to better characterize the pathophysiology in patients with *Gata2* germline mutation.

## Methods

### Mice

C57BL/6N mice were purchased from Janvier (Orleans, France) and CD45.1 B6.SJL congenic mice from Jackson Laboratory (Bar Harbor, ME, USA). *Gata2*^+/-^ mice were kindly provided by Pr. Stuart Orkin (Harvard Stem Cell Institute, USA) and backcrossed to C57BL/6N background. *Gata2^R396Q/+^* mice was designed by us and generated by the CIPHE animal facility (Marseille, France) using homologous recombination into ES cells and a homologous recombination vector containing from 5’ to 3’, a 2732 base pairs (bp) long 5’ homology arm (from the beginning of exon 4 to the end of exon 5), a loxP site, the wildtype exon 6 tagged by a V5 sequence, a stop codon, a LoxP511 sequence and in a reverse orientation, a stop codon, an OST (One Strep Tag) sequence, the exon 6 containing the R396Q mutation, a loxP and a Lox511 sequences, a FRT-NeoR-FRT cassette and a 3000 bp 3’ homology arm of the 3’UTR. Chimera mice were breed with C57Bl6/N mice and recombinant offspring were bred with Flp deletor mice to remove the FRT-NeoR-FRT cassette. These mice bred with Vav-Cre transgenic mice leading to the selection of mice containing the mutated exon 6 in the correct orientation (G*ata2^R396Q/+^* mice). All experiments were conducted according to French law and approved by the local ethics committee for animal experimentation (Protocol numbers: 19-16U1037-CB-01 and 20-16U1037-CB-02).

### Genotyping

For genotyping PCR, DNA was extracted from mice ear tissues (or tail for embryos) using Extraction « Extract-N-Amp PCR » kit (Sigma #XNAT2-1KT) according to the manufacturer’s instructions. The PCR reaction was performed on a Veriti thermocycler (Applied Biosystems) with Gata2-forward (5’ AGGCTGTGCAGGCATTTA 3’) and Gata2-reverse (5’ CTGCCAAACCACCCTTGAT 3’) primers using the following conditions: 94°C for 3 min; 31 cycles of 94°C for 30 sec, 58°C for 40 sec, 72°C for 30 sec; and 72°C for 7 minutes. A 796bp (corresponding to the mutated allele) and/or a 592bp (corresponding to the wt allele) DNA bands were detected on a TapeStation DNA ScreenTape (Agilent).

For *Gata2*^+/-^ mice, the mutated allele was detected by amplified a band of 650 bp using a Neospecific PCR with neo 504 (5’-GCCCGGTTCTTTTTGTCAAGACCG-3’) and Neo505R (5’-CAGAAGAACTCGTCAAGAAGGGGA-3’) with Hotstart master Mix (Ozyme#OZYA009-20) using the following conditions: 95°C for 15 minutes; 31 cycles of 94°C for 30 sec, 60°C for 45 sec, 72°C for 1 min; and 72°C for 10 minutes.

### Clonogenic assays

A 5’ 3xHA-Tag sequence were inserted into pMSCV-IRES-GFP II (MIG) plasmid (Addgene, Plasmid #52107;23). Human GATA2 WT, GATA2 R204X and GATA2 R396Q complementary DNAs (cDNAs) were subcloned into this MIG plasmid. The Phoenix-eco retroviral packaging system/cell line (φNXe) (http://www.stanford.edu/group/nolan) was used to generate MIG GATA2 WT, MIG GATA2 R204X, MIG GATA2 R396Q and control MIG (expressing only the 3xHA-Tag sequence) ecotropic retroviruses.

Retroviral supernatants were produced by transient transfection of Phoenix cells pre-incubated in IMDM (Gibco #21980) or DMEM (Gibco #61965), one hour prior to transfection and kept along transfection. After overnight incubation, cells were treated for 6 h in a medium containing 10 mM sodium butyrate (Sigma #B5887). Viral supernatants were harvested after at least 24 h of incubation at 32°C and filtered on a 0.45 µm filter (Merck #SE2M230I04).

Murine lineage negative cells, transduction was performed using pure retroviral supernatant supplemented with 8 µg/mL polybrene (Sigma# H9268), 10ng/mL mIL3; 20 ng/mL hIL6 and 40 ng/mL mSCF. (Peprotech respectively #213-13, #200-06 and #250-03). Spinoculation was performed at 1,000*g* at 32°C for 90 min. After centrifugation, medium was replaced by IMDM (Gibco #21980) supplemented with 10% of Fetal Bovine Serum (FBS, Gibco #10270) and cytokines as above. Cells were cultured at 37°C with 5% CO_2_ in StemSpan (Stem Cell Technology #09650) supplemented with TPO (0,125ng/mL), FLT3L (1ng/mL), SCF (0,25ng/mL) and 10% of SVF during 3 days after transduction, then GFP^+^ cells were sorted using BD FACSAria™ III Cell Sorter (BD Biosciences)

Transduced lineage negative (Lin^-^) or Lin^-^Sca-1^+^Kit^+^ (LSK) cells were plated into 1mL of methylcellulose-based medium with cytokines (Stem Cell Technology, MethoCult GF M3434) into 35 x 10 mm non-adherent Bio-One Petri Dish, 35 x 10 mm, (Greiner # 627102) and placed into a wet chamber at 37°C and 5% CO2. Colonies were quantified on a Leica microscope at x20 magnification. After incubation, cells were recovered by dilution in 37°C pre-warmed PBS 1X, washed twice into PBS 1X and 10,000 cells were plated into a 1mL fresh MethoCult medium for serial passages. In parallel, 100,000 cells were cytocentrifuged at 500 rpm for 10 min and the remaining cells were analyzed by flow cytometry.

### Immunofluorescence

#### BAF/3 cell lines

Viral supernatants were diluted 1/3 in RPMI supplemented with 10% FBS, Sodium Pyruvate (Sigma #S8636), 56µM of beta-mercaptoethanol (Sigma #3148), 8 µg/mL polybrene (Sigma #H9268), 10ng/mL mIL3 (Peprotech #213-13). Spinoculation was performed at 1,000*g* at 32°C for 90 min. After centrifugation, medium was replaced by BaF/3 medium. Cells were incubated overnight at 37°C and 5% CO_2_ and sorted by GFP expression with an Aria III cell sorter (Becton Dickinson).

Sorted cells were seeded on poly-L lysine (Sigma #P8920-100mL), then fixed with 4% PFA (Electron Microscopy Sciences #15713) for 15 min and washed 3 times in PBS 1X then saturated in PBS 1x with 5% BSA for 30 min. Cells were permeabilized with triton-X100 (Sigma #T8787). Cells were stained for 1h at RT with a monoclonal Anti-HA tag antibody [16B12] (ab130275, 1/100) and a secondary goat anti-mouse antibody labeled with Alexa 546 (Thermofisher #A-21123, 1/500) then counterstained with Prolong Gold mounting medium containing DAPI (Life Technologies #P36941).

Immunofluorescence was revealed on an AXIO microscope (ZEISS), and quantification was performed on 10 HA positive cells (except for the control condition on GFP positive cells) using ZEN software (Zeiss).

#### E12.5 Fetal liver

Whole fetal livers (FL) were fixed during 24h at 4°C in 4% formaldehyde and then embedded in 4% low-melting agarose (Sigma #A9539-500G). One hundred µm sections were made using a vibratome (Leica #VT1200) as described here^23^. Tissues were blocked and permeabilized with buffer containing 10% FBS, 20% DMSO and 0.1% Triton X-100, incubated with a fluorescently labelled antibody Endomucin (Rat anti-mouse, clone TC15-12F12.2, BioLegend) at 4°C overnight under agitation. Immunostaining with CF633-labelled donkey anti-rat antibody was added the next day for 4 hours and washed. FL sections were incubated with a labelled CD150 (PE, Rat anti-mouse) antibody at 4°C overnight under agitation. The fluorescently labelled tissues were placed cut-face-down onto a microslide and were covered in Dako mounting medium (Dako#S302380) to prevent tissue desiccation. The preparations were examined under LSM780 and LSM 880 (Zeiss) confocal microscopes and analyzed with Zen software (Zeiss).

### Flow cytometry, cell sorting and data analyses

Single cell suspensions were prepared from bone marrow (BM) mice of IMDM supplemented with 2% FBS. Immunostainings were performed using antibodies for flow cytometry obtained from Pharmingen (BD Biosciences), Invitrogen (Thermo Fisher Scientific) and Miltenyi Biotec. Flow cytometry experiments were performed with BD antibodies (listed as supplemental information). For surface staining, cells were incubated with the antibodies for 20 minutes in IMDM 2% FBS at 4°C and washed twice with PBS 1X before analysis. For cell-cycle analysis and intra-cellular staining, cells were fixed and permeabilized (Transcription Factor Fix/Perm Buffer, BD Bioscience #562725) overnight at 4°C before the staining with the anti-Ki67, anti-Gata2, anti-Gata1 antibodies in PermWash Buffer. Flow cytometry experiments were performed using a BD LSRFortessa™ X-20 Cell Analyzer (BD Biosciences) using DIVA software (BD Biosciences).

Prior to cell sorting, BM cells from *Gata2*^+/+^ or *Gata2*^R396Q/+^ mice were flushed, and lineage negative cell fraction were enriched using EasySep Mouse Hematopoietic Progenitor Cell Isolation Kit (Stem Cell Technologies #19856). Cell sorting of LSK or LT-HSC, ST-HSC and MPP3-4 was performed using a BD FACSAria™ III Cell Sorter (BD Biosciences) using DIVA software (BD Biosciences).

flow cytometry data were analyzed using FlowJo Version 10.8 software (BD Life Sciences). Data cleansing and normalization were performed using FlowAI package (V2.1)^24^ for each sample, and the same number of cells were then concatenated. Arc sin transformation was performed manually for each marker to discriminate the different LSK subpopulations. The dimension reduction algorithm TriMap^25^ was run. TriMap analyses were performed using our multiparametric staining covering all LSK subpopulations. In dimensional reduction representation, each subpopulation is represented by one color.

### Transcriptomic analysis

#### LSK

Total RNA from 60,000 purified LSK *Gata2*^+/+^ and *Gata2*^R396Q/+^ mice was isolated with the Rneasy Plus Mini Kit (Qiagen, #74134). RNA-seq libraries were prepared using TruSeq Stranded mRNA Low Sample kit (Illumina) according to manufacturer’s protocol, starting with 300 ng of total RNA. Cluster generation and sequencing were carried out using the Illumina NextSeq 500/550 High Output Kit v2.5 (150 cycles) with a read length of 2 x 75 nucleotides.

#### LT-HSC, ST-HSC, MPP3-4

Specific LSK subpopulations from *Gata2*^+/+^ and *Gata2*^R396Q/+^ mice were purified by cell sorting (BD FACS Melody™ Cell Sorter (BD Biosciences) directly in RLT buffer, and total RNA was extracted with the RNeasy Plus Micro Kit (Qiagen, #74034). Smart-seqv4 libraries were prepared as previously described^26^ using the Takara SMART-Seq v4 full-length transcriptome analysis kit. Paired-end sequencing was performed on an Illumina NextSeq 500 using 2 x 75 bp reads.

Low quality reads were trimmed, and adaptor sequence was removed using fastp (v0.23.2). Short reads were then mapped to mm10 genome using RNA-STAR (v2.7.8a) with doublepass parameter. Low quality mapping and duplicated reads were removed, and the remaining reads was count using featurecount (v2.0.1) with default paired-end parameter.

### ATAC-Seq

Sorted LSK populations from *Gata2*^+/+^ and *Gata2*^R396Q/+^ mice were prepared according to the Active-Motif ATAC-Seq kit (#53150)^27^. The size distribution and concentration of the libraries were assessed on Tapestation with a DNA High Sensitivity kit (# 5067-4626, Agilent Technologies, Santa Clara, CA). Paired end 38 bp sequences were generated from samples on a NextSeq 500/550 (Illumina, San Diego, CA, USA) with an average of 125 million of reads per sample.

First, FastQC is used to assess the sequence quality. Foreign sequences removal and trimming are realized with Sickle (qual threshold 20 and length threshold 20). Sequences were mapped to the murine genome with Bowtie2 (2.4.2)^28^ with -X 2000 (maximal fragment length), very-sensitive and against mm10. Pipelines were used for various cleaning and filtering steps^29^ : removing mitochondrial reads and reads from non-assembled contigs or alternative haplotypes, filtering reads with mapping quality <20, marking and removing duplicate reads and adjusting read start sites as described previously^27^ (4 bp on the forward and 5 bp on the reverse strand). Lastly, a GC bias diagnosis and correction using *deepTools* was run for each sample. The output of this pre-processing pipeline was used to peak calling using MACS2 with -SE -200 -100 - lambdafix parameter and removal of blacklisted regions.

#### DiffTF analysis

The complete diffTF pipeline^29^ was run using TF binding sites (TFBS) generated by PWMscan analysis (cutoff p-val: -0.00001, background base composition: - 0.29;0.21;0.21;0.29) of each JASPAR2020 PWMs motifs to obtain position of non-redundant and specific motifs. The analytical approach was preferred due to the small size of the sample and paired design was used for DESeq2. For the plotting of individual activity for each TF, we used HINT-ATAC pipeline^30^ with standard parameter on the combination of the bam of each condition for the JASPAR2020 database.

### Functional *in vivo* assays

For BM reconstitution assays, myeloablation was induced by two doses (250mg/kg) of 5-Fluorouracil (Accord) administered intraperitoneally at days 0 and 11.

Acute inflammatory stress was performed using lipopolysaccharide (Sigma #L2630) injected intraperitoneally at 5 mg/kg in 8-week-old mice. Mice were euthanized after 16 hours or 96 hours. Murine bone marrow cells were characterized by flow cytometry.

Transplantations of LSK cells were performed after a myeloablative conditioning using a 30 mg/kg sublethal dose of busulfan (Sigma #B2635) injected into the peritoneal cavity of 8-week-old recipient mice 24 hours before intravenous transplantation. Cell chimerism was quantified by flow cytometry as the percentage of donor-derived cells (% of CD45.2^+^ cells in total CD45^+^ BM cells) found in the CD45.1^+^ recipient bone marrow.

Transplantation of LT-HSCs was performed after lethal irradiation (9 Gy) 6 hours before the transplantation. Five hundred LT-HSCs from CD45.2^+^ *Gata2*^+/+^ or *Gata2*^R396Q/+^ (CD150^high^ or CD150^low^) mice with one million of total bone marrow CD45.1^+^ cells as support were injected in CD45.1^+^ recipient mice.

### WGS and RNA-Seq

#### DNA Extraction

Constitutional genomic DNA is extracted from total bone marrow at the remission and purified using magnetic beads (Maxwell® RSC Whole Blood DNA Kit, PROMEGA). Tumor genomic DNA is extracted and purified using magnetic beads (Maxwell® RSC Tissue DNA Kit, PROMEGA), either through automated means (Maxwell® RSC System, PROMEGA) or manually (AllPrep DNA/RNA Mini Kit, QIAGEN).

#### RNA Extraction

Tumor RNA is extracted and purified using magnetic beads (Maxwell® RSC Simply RNA tissue Kit, PROMEGA), either through automated means (Maxwell® RSC System, PROMEGA) or manually (AllPrep DNA/RNA Mini Kit, QIAGEN).

#### WGS Library Preparation

Fragmentation is obtained through sonication (Covaris L220plus® COVARIS®). Library preparation is performed with or without amplification (Nextera® DNA Flex Library Prep kit or TruSeq® DNA PCR-Free kit, ILLUMINA®), either through automation (MICROLAB Star®, HAMILTON®) or manually. Size profiles are analyzed through capillary electrophoresis (TapeStation 4200®, Agilent®). The library is quantified through qPCR (Invitrogen™ Collibri™ Library Quantification Kit; QuantStudio™ 5 Real Time PCR Systems, APPLIED BIOSYSTEMS).

#### RNA-seq Library Preparation

The RNA depletion step is performed using magnetic beads (Agencourt® RNAClean XP, BECKMAN COULTER), while the other purification and size selection steps are performed using magnetic beads (Agencourt® AMPure XP, BECKMAN COULTER). Library preparation from RNA is performed with amplification (TruSeq Stranded Total RNA H/M/R-Gold, ILLUMINA®), either through automation (MICROLAB Star®, Hamilton®) or manually. Size profiles are analyzed through capillary electrophoresis (TapeStation 4200®, Agilent®). The library is quantified through spectrofluorometric measurement (Quant-It™ kit, INVITROGEN™, Spark™, TECAN®).

#### Library Sequencing

The libraries are sequenced in paired-end (2 x 150 cycles) through the SBS (sequencing by synthesis) technology using NovaSeq® 6000 (ILLUMINA®).

#### Alignment of Normal and Tumor Genomes (DNA)

Demultiplexing is performed using Illumina’s proprietary bcl2fastq-v2-20-0 tool. Raw sequence data from normal and tumor genomes are then aligned to the GRCh38.p13 version of the human reference genome using the bwa-mem-v0.7.17 tool. Duplication marking is performed using biobambam-v2.0.89. BAM files was visualized using IGV (Integrated Genomics Viewer).

### Statistics

Statistical differences were determined using a two-tailed unpaired Student’s t test for comparison of quantitative variables, when assuming normality and equal distribution of variance between the different groups analyzed otherwise using Mann-Whitney test. Error bars for pooled replicates represent standard deviation (SD) unless specified (SEM, standard error of the mean). Survival differences were analyzed using log-rank test. A *p* value lower than 0.05 was considered as statistically significant (*p < 0.05; **p < 0.01; ***p < 0.001; ****p< 0.0001). All statistical analyses were performed using GraphPad Prism software, version 7 (GraphPad).

## Results

### Ectopic expression of GATA2^R396Q^ induces an excessive granulocytic differentiation *in vitro*

To date, 179 different *GATA2* pathogenic or likely pathogenic germline variants have been identified in patients^13^. Nonsense and frameshift coding mutations^6, 8, 10, 12, 16, 31–36^ (Table S1) have been consistently reported upstream to the second zinc finger (C-ZF) (Figure 1A) leading to an haploinsufficiency^14^. In contrast, *GATA2* germline missense mutations are clustered on the second C-ZF and associated with a higher risk of leukemic transformation^16, 37^. In the C-ZF, germline mutations recurrently impact arginines (R361, R362, R396 and R398)^13^. Among them, mutation affecting arginine 396 is considered as a mutational hotspot. We then investigate the impact of the variant GATA2^R396Q^ on the hematopoietic system.

**Figure 1.**
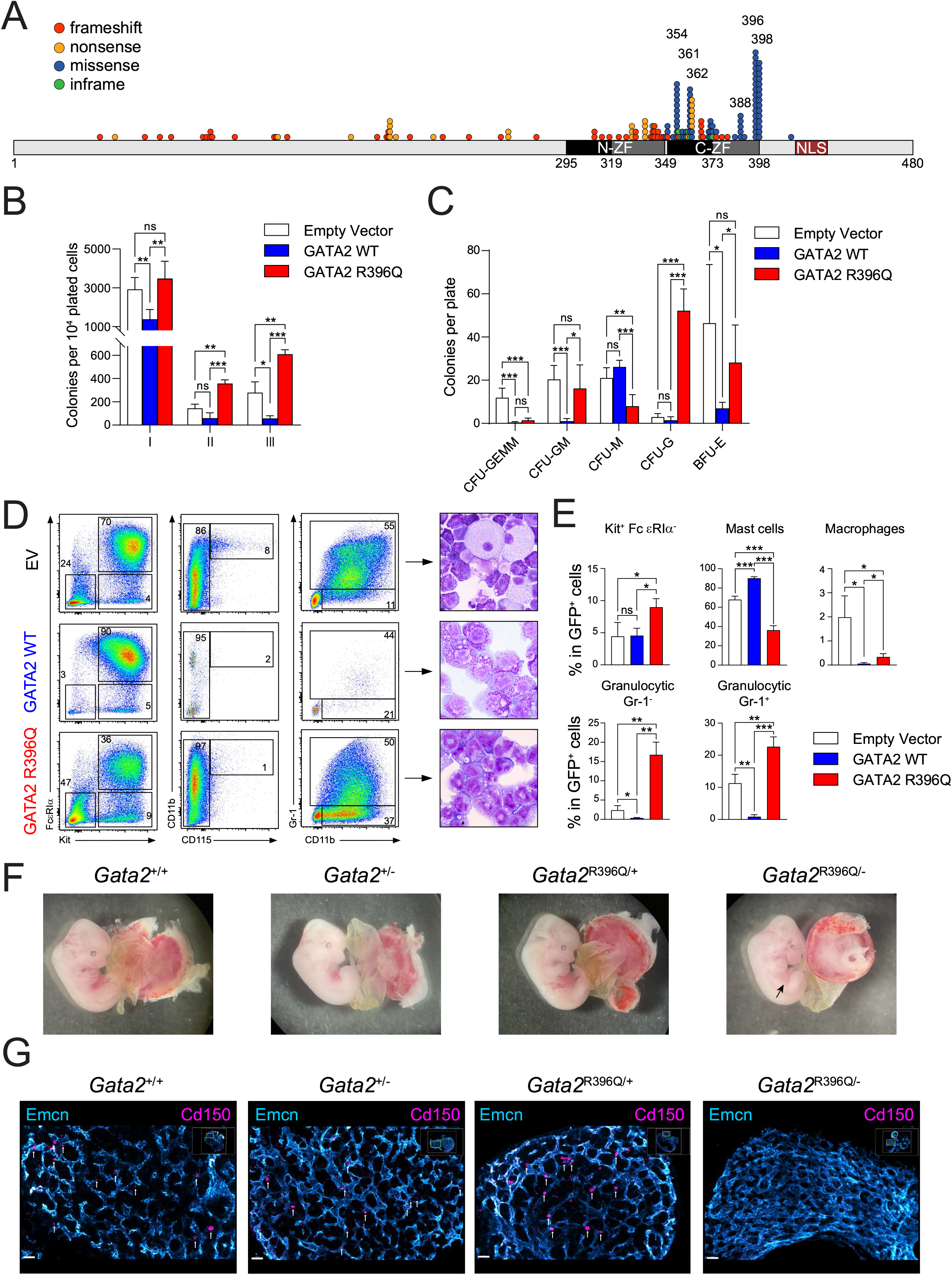
The recurrent *Gata2*^R396Q^ mutation impairs the emergence and the differentiation of hematopoietic cells. (A) Compilation of *GATA2* germline mutation cases previously reported in literature^6, 8, 10, 12, 16, 31–36^. Each dot represents a patient with alteration types color-coded. (B-E) Clonogenic assay performed on murine lineage negative (Lin^-^) cells transduced with viruses encoding GFP alone (Empty vector), GFP and GATA2 wild-type (GATA2 WT), or GFP and GATA2 R396Q (GATA2 R396Q). GFP^+^-sorted Lin^-^ cells were plated on cytokine-supplemented methylcellulose medium. (B) Quantification of colonies, with GFP^+^ cells serially plated every 7 days (n=6). (C) Identification and quantification of colony types 10 days after the initial cell plating. (D) Immunophenotyping by flow cytometry (left panel) and May-Grunwald Giemsa coloration (right panel) of cells recovered from methylcellulose after the third cell plating (n=3). (E) Statistical results of FACS analyses of cells recovered after the third cell plating. (F) Images of E12.5 embryos resulting from *Gata2^+/-^*x *Gata2*^R396Q/+^ mating. (G) Immunostaining of fetal liver sections, with Endomucin (Emcn) in blue and CD150 in purple. Scale bar represents 40 µm. ZF: Zinc finger domain; NLS: nuclear localization signal; CFU: Colony forming unit; GEMM: Granulocyte, Erythroid, Macrophage, Megakaryocyte; GM: Granulocyte, Macrophage; M: Macrophage; G: Granulocytes and BFU-E: Burst-forming unit erythroid. Results are shown as mean ± SD; ns: not significant, *p<0.05, **p<0.01, ***p<0.001, ****p<0.0001.

We first confirmed, by ectopic expression on BaF/3 cells, the capacity of this variant to maintain a nuclear localization similarly than GATA2^WT^, while GATA2^R204^*, a truncated variant missing the NLS domain, remained in cytoplasmic compartment (Figure S1A-B).

The *GATA2*^R396Q^ mutation was reported to have a significantly reduced DNA-binding and transactivation capacities^37^. To assess the functional consequences of this missense mutation, we transduced murine lineage-negative (Lin^-^) cells with a GFP-expressing virus encoding either GATA2^WT^ or GATA2^R396Q^ (Figure S1C-D). Ectopic expression of GATA2^WT^ significantly reduced the number of colonies and induces mast cell differentiation (Kit^+^ FcεRIα^+^) consistent with the reported inhibitory effect of GATA2 overexpression on normal hematopoiesis^38^ (Figure 1B and 1D-E). Conversely, GATA2^R396Q^ enhanced clonogenic capacity compared with both empty vector and GATA2^WT^ (Figure 1B) and induced a differentiation towards granulocytic differentiation, with an increase in number of CFU-G colonies (Figure 1C) and in proportion of immature and mature granulocytes (CD11b^+^ Gr-1^+/-^) (Figure 1D-E and S1D). These results demonstrated that GATA2^R396Q^ is not equivalent to a null mutation or to GATA2^WT^, as reported for other missense GATA2 variants impacting N-ZF and C-ZF domains^15, 37^ suggesting ectopic functions of this mutant.

### Gata2^R396Q^ is ineffective in initiating early hematopoietic stem cell development

To mimic patients molecular context, we generated a *Gata2*^R396Q/+^ knock-in mouse (Figure S1E-F). We first analysed the progeny of the breeding of a *Gata2*^R396Q/+^ mouse with a *Gata2^+/-^*mouse at E12.5, just after the yolk sac hematopoietic waves^39^ and detected all genotypes in the expected proportions (Figure S1H and Table S2). However, in contrast to all the other embryos, *Gata2*^R396Q/-^ embryos displayed gross developmental abnormalities including a translucent fetal liver (Figure 1F and S1G)) with a lack of HSCs as demonstrated by the absence of CD150^+^ cells (Figure 1G). Expectedly, no *Gata2*^R396Q/-^ mouse were obtained at birth (Table S2, *p*=0.012), reminiscent of the *Gata2*^-/-^ mice^40^. Therefore, Gata2^R396Q^ was unable to rescue the phenotype associated with the loss of the wild-type allele, suggesting that Gata2 functions, crucial for the early development of hematopoietic cells, are impaired due to the mutation.

### *Gata2*^R396Q^ mutation disrupts LSK distribution throughout mouse development and aging

We further analyzed *Gata2*^R396Q/+^ hematopoietic development. At E14.5, the number of LSK cells in the fetal liver is increased in *Gata2*^R396Q/+^ compared with *Gata2*^+/+^ embryos, mainly attributed to a significant rise in the number of Long-term (LT)-HSCs and Multipotent Progenitors (MPP)2 cells, while the count of Short-term (ST)-HSC cells decreased (Figure 2A and S2A). Furthermore, in 2-month-old mice, this altered distribution of subpopulations observed during embryonic development persisted, accompanied by an additional increase in MPP3 cells (Figure 2B and S2D). Cell cycle analysis using Ki67 staining, did not reveal significant differences, except in *Gata2*^R396Q/+^ MPP2 cells, where a lower proportion was found to be cycling (Figure S2C). This suggests that the observed variations in cell numbers between *Gata2*^R396Q/+^ and wild-type mice were not mainly due to differences in cell cycle activity.

**Figure 2.**
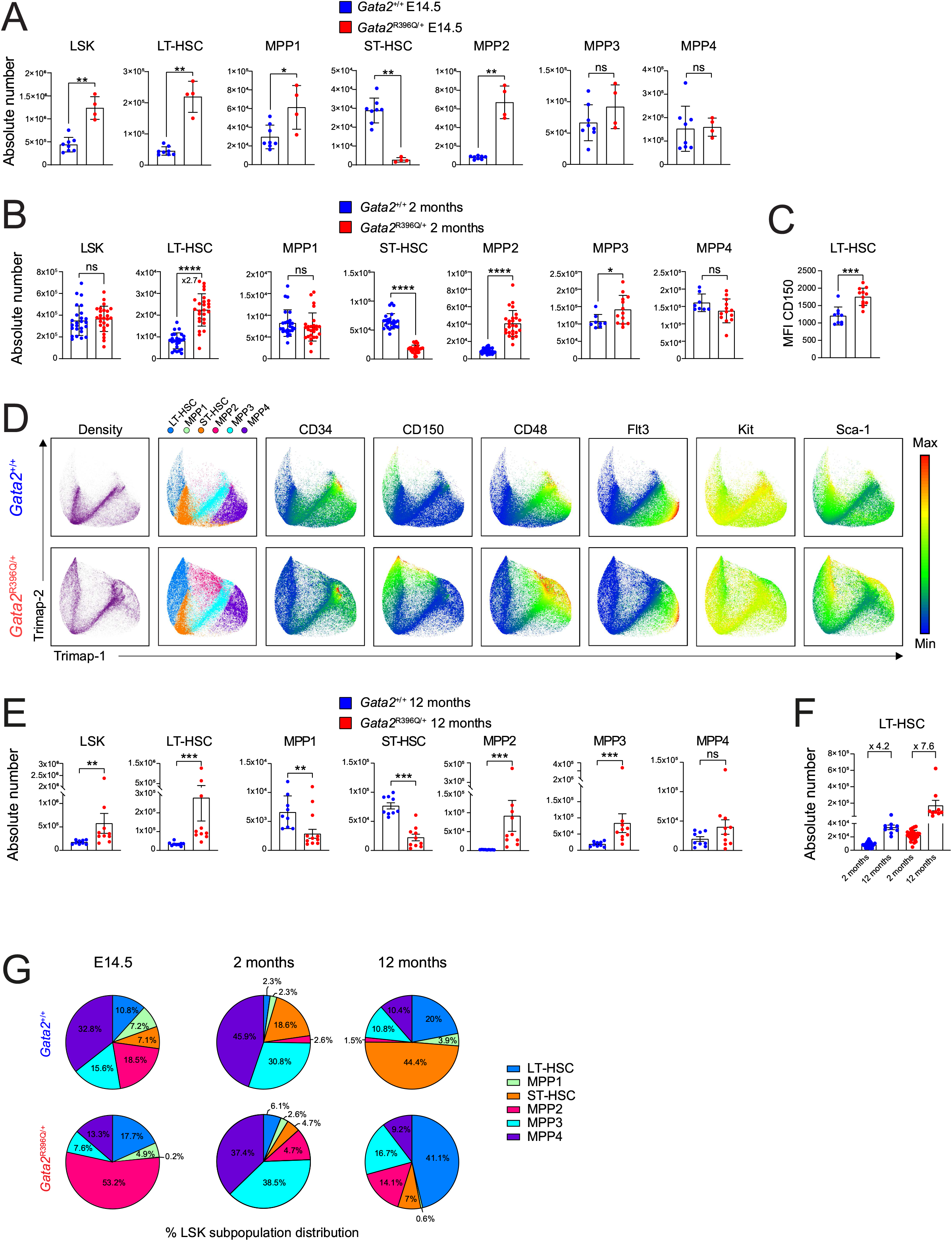
Repartition of LSK subpopulations in Gata2^R396Q/+^ mice from the embryonic stage to middle age. (A) Absolute number of LSK, LT-HSC, MPP1, ST-HSC, MPP2, MPP3, and MPP4 cells in E14.5 fetal liver of *Gata2^+/+^* (blue, n=8) and *Gata2*^R396/+^ (red, n=4) embryos. (B) Absolute number of LSK, LT-HSC, MPP1, ST-HSC, MPP2 (n=25), MPP3 and MPP4 (n=8 *Gata2*^+/+^, n=13 *Gata2*^R396Q/+^) cells in 2-month-old *Gata2^+/+^* (blue) and *Gata2*^R396/+^ (red) mice. (C) Mean fluorescence intensity (MFI) of CD150 marker on LT-HSC surface of 2-month-old *Gata2^+/+^* (blue) and *Gata2*^R396/+^ (red) mice. (D) TriMap representation of LSK subpopulations from 2-month-old mice, showing population density and marker intensity (CD34, CD150, CD48, Flt3, Kit, and Sca-1). A colored scale indicates marker intensity. (E) Absolute number of LSK, LT-HSC, MPP1, ST-HSC, MPP2, MPP3, and MPP4 cells in 2-month-old *Gata2^+/+^* (blue, n=9) and *Gata2^R396/+^*(red, n=10) mice. (F) Comparison of the absolute number of LT-HSC in the bone marrow of 2- and 12-month-old *Gata2^+/+^* (blue, n=9) and *Gata2*^R396/+^ (red, n=10) mice. Fold increase between 2 and 12 months is indicated. (G) Distribution (in percentage) of each LSK subpopulation in E14.5 embryos, 2- and 12-month-old *Gata2^+/+^*and *Gata2^R396/+^* mice. LT: Long term; ST: Short term, HSC: Hematopoietic stem cell, MPP: Multipotent progenitor. Each dot represents an individual mouse. Results are shown as mean ± SD; ns: not significant, *p<0.05, **p<0.01, ***p<0.001, ****p<0.0001.

Our phenotypic characterization revealed significant disparities with the heterozygous *Gata2*^+/-^ model mainly characterized by a reduction in the number of LT-HSCs (Figure S2F). Furthermore, we highlighted a population of mutated LT-HSCs, characterized by higher expression of CD150 and Sca-1 markers, and lower expression of Kit compared with *Gata2*^+/+^ and *Gata2*^+/-^ LT-HSCs (Figure 2C-D and S2B, E and G).

At one year-old, defined as middle age, LSK cell number was higher in *Gata2*^R396Q/+^ mice than in *Gata2*^+/+^ mice (Figure 2E). Physiologically, aging results in an increased proportion of CD150^high^ LT-HSCs associated with a myeloid-biased differentiation pattern^41, 42^. Interestingly, in *Gata2*^R396Q/+^ mice, the LT-HSC population rise with a higher fold from 2 to 12 months of age when compared with wild-type mice (7.6-*versus* 4.2-fold) (Figure 2F). It is worth noting that LSK subpopulation misdistribution was a constant hallmark throughout development and life of *Gata2*^R396Q/+^ mice (Figure 2G).

Collectively, these findings demonstrate that Gata2^R396Q^ exhibits a distinct ectopic function, which goes beyond a simple loss of function, significantly impacting the hematopoietic stem cell compartment.

### Alteration of stem cells affects the mature compartment

Bone marrow characterization of committed Lin^-^Kit^+^Sca-1^-^ (LK) progenitor numbers of 2-month-old *Gata2*^R396Q/+^ mice revealed a decrease compared with their wild-type counterparts mainly due to a reduction in CMP cells and, to a lesser extent, MEP cells (Figure 3A and S3A-C). In *Gata2^+/-^* mice of the same age, no difference in LK progenitor numbers was observed compared with wild-type mice (Figure S3D). The absolute numbers and proportions of mature cells including myeloid, B and T cells in the bone marrow were similar (Figure 3B and S3E). However, in the blood, the hemoglobin level and platelet count were decreased in mutated mice (Figure 3C). Similar analysis performed on *Gata2*^+/-^ mice and their wild-type counterparts showed no difference in mature cells (Figure S3D,F), reinforcing the notion of an aberrant function attributed by this missense mutation compared to a loss of one functional allele.

**Figure 3.**
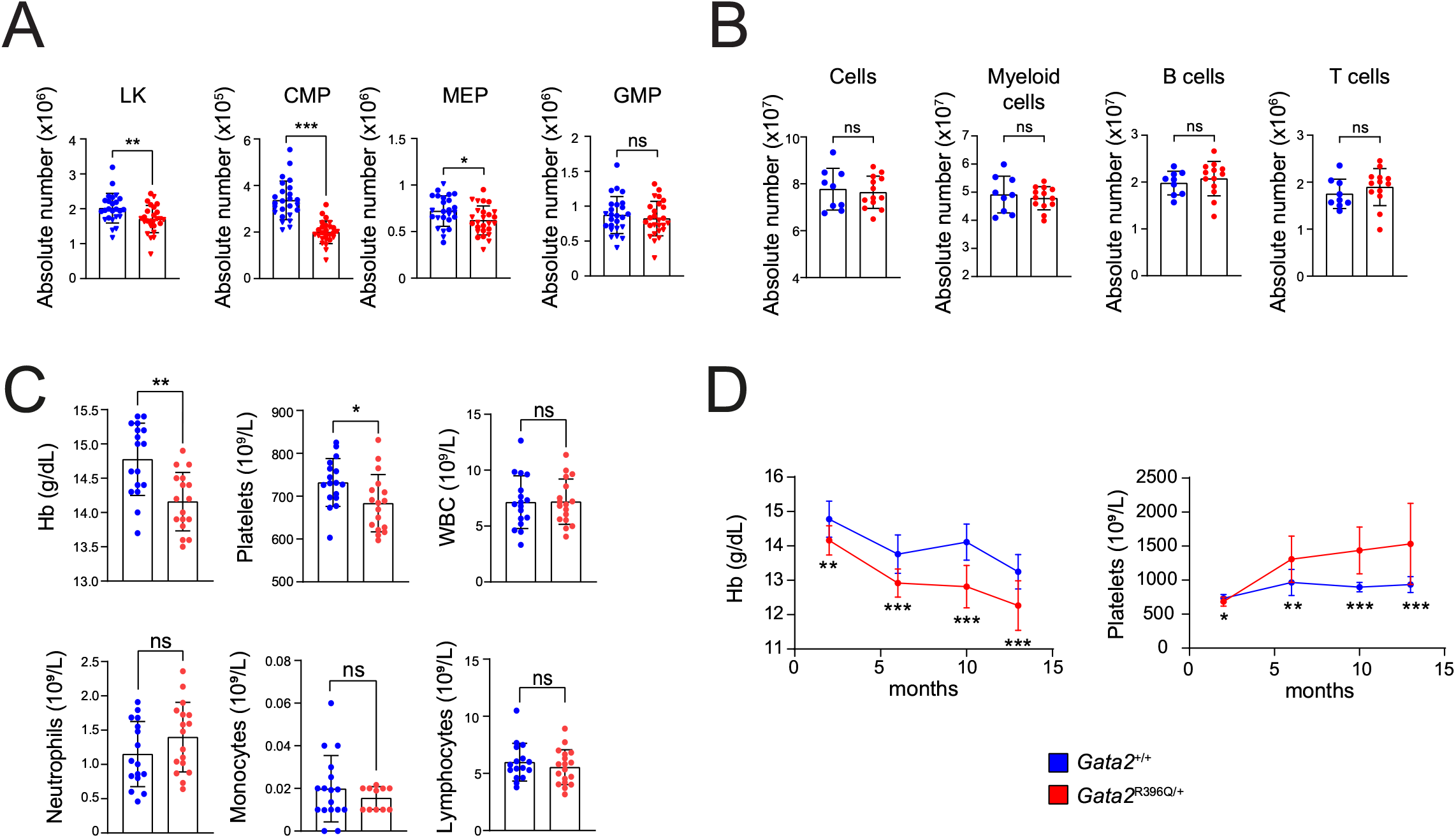
Impact of the mutant on early and late hematopoiesis. (A) FACS analysis statistics of the absolute number of LK, CMP, GMP, and MEP cells in the bone marrow of 2-month-old *Gata2^+/+^* (blue, n=25) and *Gata2*^R396/+^ (red, n=25) mice. (B) FACS analysis statistics of the absolute number of total cells, Myeloid cells, B cells, and T cells in the bone marrow of 2-month-old *Gata2^+/+^* (blue, n=9) and *Gata2*^R396/+^ (red, n=13) mice. (C) Blood parameters, including hemoglobin level (Hb) and white blood cells (WBC), in 2-month-old *Gata2^+/+^* (blue, n=17) and *Gata2*^R396/+^ (red, n=11) mice. (D) Time course of hemoglobin level and platelet number in the blood of *Gata2^+/+^* (blue) and *Gata2*^R396/+^ (red) mice. Hb: Hemoglobin level, WBC: white blood cells. LK: Lineage^-^ Kit^+^ cells; CMP: Common myeloid progenitor, MEP: Megakaryocyte–erythroid progenitor, GMP: Granulocyte-monocyte progenitor. Each dot represents an individual mouse. Results are shown as mean ± SD (except (E), SEM); ns: not significant, *p<0.05, **p<0.01, ***p<0.001, ****p<0.0001.

Time tracking of blood parameters showed that *Gata2*^R396Q/+^ mice develop anemia over time while platelet number increase rapidly, indicating differentiation bias in favor of platelet over erythroid development (Figure 3F). Additionally, middle-aged mice exhibited an elevated number of granulocytes in the blood (Figure S3G). Altogether, we demonstrated that quantitative defects in the stem cell compartment affect terminal hematopoietic differentiation, with a greater impact on the megakaryocytic and erythroid lineages. The effects on this mature compartment became increasingly pronounced as the mice age. Aging is a physiological process that can be viewed as a challenge for hematological compartments. Therefore we aimed to investigate whether the functionality of immature cells was affected, when they were challenged at an early age.

### *Gata2*^R396Q^ mutation impairs LSK development and function

To assess the intrinsic differentiation potential of *Gata2*^R396Q/+^ compared to *Gata2*^+/+^ LSK cells from 2-month-old mice, we conducted clonogenic assay. *Gata2*^R396Q/+^ cells exhibited a significant decrease in CFU-GEMM colonies (Figure 4A), and when subjected to serial passages, these cells displayed an increase in colony number (Figure S4A). Morphological and phenotypic analyses revealed a predominance of granular precursors and mature granulocytes in the remaining *Gata2*^R396Q/+^cells whereas mast cells were more predominant in *Gata2*^+/+^ condition (Figure S4B-C). *Gata2*^+/-^ LSK cells displayed reduced CFU-GEMM and CFU-GM without showing increased clonogenic activity. As for *Gata2*^+/+^, mast cells were the most prominent cells observed in *Gata2*^+/-^ condition (Figure S4D-F).

**Figure 4.**
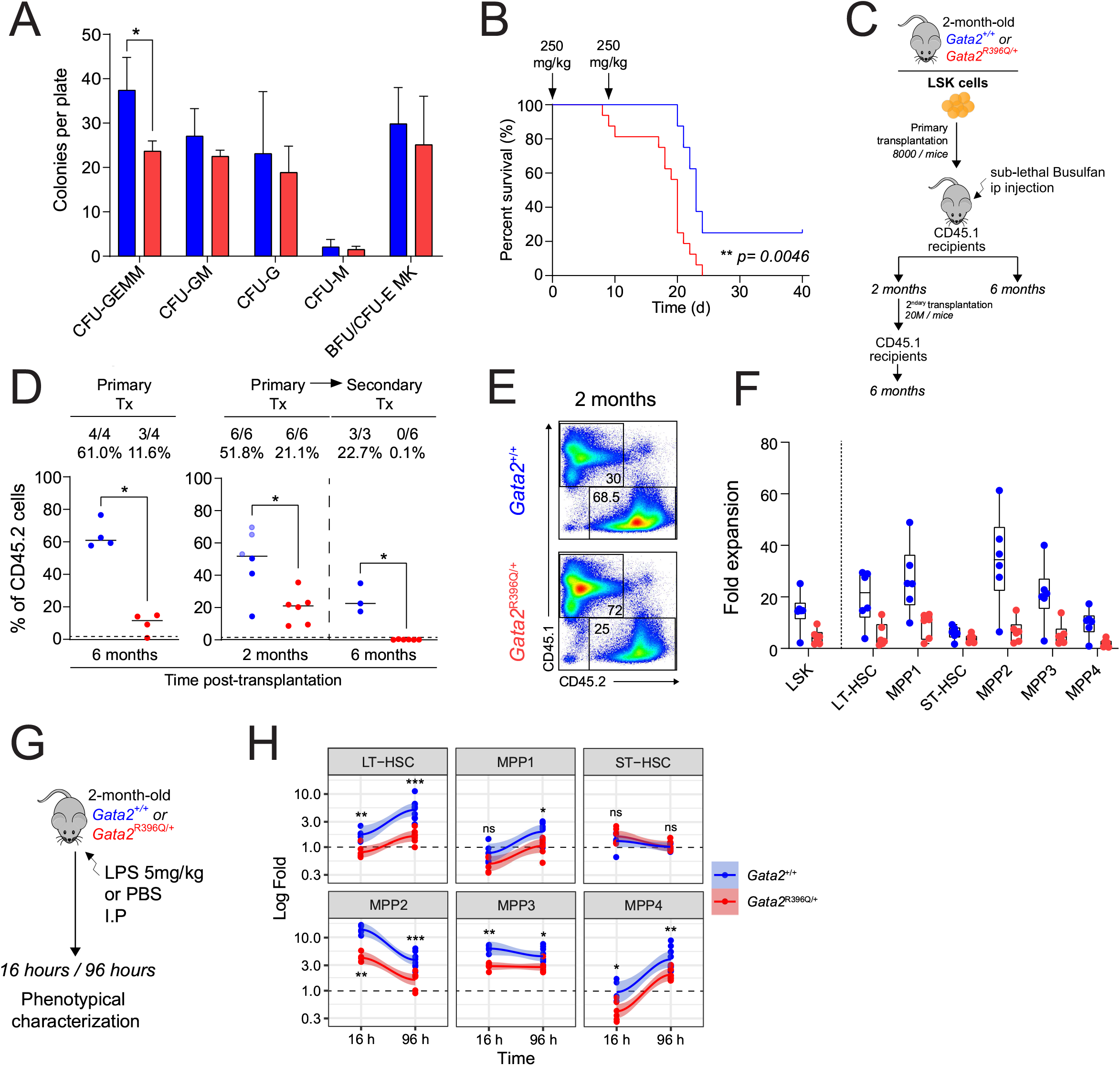
*Gata2*^R396Q/+^ mice exhibit qualitative defect in challenging conditions. (A) Clonogenic assay on 2-month-old *Gata2*^+/+^ (blue) and *Gata2*^R396Q/+^ (red) Lin^-^Sca-1^+^Kit^+^ (LSK) cells. Colonies were counted after 10 days (3 independent methylcellulose assays). (B) Kaplan-Meier survival curve (in days) of *Gata2*^+/+^ (blue, n=8) and *Gata2*^R396Q/+^ (red, n=16) mice after two injections (arrows) of 5-fluorouracil (5-FU) (C) Experimental scheme for transplantation assays. (D) Engraftment quantification of CD45.2^+^ cells (from donor mice) 6 (left panel) and 2 (left part of the right panel) months after transplantation. The number of engrafted mice (<1% of CD45.2^+^ cells) out of the total number of transplanted mice and the average reconstitution percentage for each group are indicated on top of the graph. The far-right panel shows the percentage of CD45.2^+^ cells 6 months after a secondary transplant of cells from mice indicated with dark-colored dots. Each blue dot represents an individual *Gata2*^+/+^ mouse, and red dots represent *Gata2*^R396Q/+^ mice. (E) Representative flow cytometry dot plot showing bone marrow engrafted cells (CD45.2+) from *Gata2*^+/+^ and *Gata2*^R396Q/+^ mice *versus* host cells (CD45.1+) two months after the first engraftment. (F) Fold expansion between the number of estimated engrafted cells and the number recovered 2 months after engraftment for *Gata2*^+/+^ (blue) and *Gata2*^R396Q/+^ (red) cells. (G) Experimental scheme of LPS injection assay. (H) Fold expansion of specific bone marrow LSK subpopulations in *Gata2*^+/+^ (blue) and *Gata2*^R396Q/+^ (red) mice under LPS. The fold expansion is calculated as the absolute number of each LSK subpopulation in each mouse under LPS divided by the average of the absolute numbers of each subpopulation in mice under PBS. CFU: Colony forming unit; GEMM: Granulocyte, erythroid, macrophage, megakaryocyte; GM: Granulocyte, macrophage; M: Macrophage; G: Granulocytes and BFU-E: Burst-forming unit erythrocytes. E: erythrocyte, Mk: megakaryocyte. LPS: Lipopolysaccharide. Results are shown as mean ± SD; ns: not significant, *p<0.05, **p<0.01, ***p<0.001, ****p<0.0001.

To further challenge LSK cells, we assessed the BM repopulation capacity by inducing a myeloablation with 5-FU. *Gata2*^R396Q/+^ mice exhibited lower ability to reconstitute hematopoiesis, leading to a significant higher rate of death compared with *Gata2*^+/+^ mice (Figure 4B). Additionally, compared to their *Gata2*^+/+^ counterpart in a syngeneic transplantation assay, the *Gata2*^R396Q/+^ LSK repopulation capacity was found to be reduced after 2 months (Figure 4C-E) with all subpopulations showing a lower expansion (Figure 4F) except for the ST-HSCs which also did not expand in the *Gata2*^+/+^ condition (Figure S5D-E). The engraftment deficit became more pronounced after 6 months, indicating a severe impairment of *Gata2*^R396Q/+^ LT-HSCs (Figure 4D). Secondary transplantation of total bone marrow from the primary transplanted mice confirmed the loss of long-term engraftment capacity of *Gata2*^R396Q/+^ cells (Figure 4D). Even if transplantation of *Gata2^+/-^* total bone marrow cells resulted in a decreased engraftment capacity, it unveiled an imbalance in the myeloid/lymphoid ratio not seen with *Gata2*^R396Q/+^ cells. (Figure S5A-C). Overall, transplantation assays uncovered a long-term exhaustion in *Gata2*^R396Q/+^ and *Gata2*^+/-^ cells, revealed by a decrease in self-renewal capacity. Furthermore, to assess the short-term functional consequences of the *Gata2*^R396Q^ mutation on hematopoiesis, we determined the hematopoietic response after a single dose of lipopolysaccharide (LPS), mimicking an acute bacterial infection (Figure 4G). HSPCs exhibited an increased proliferation in response to inflammatory stresses^43^ (Figure S5F) triggering emergency myelopoiesis^44^. In the case of *Gata2*^+/+^ LSK cells, they expanded along with the MPP2 and MPP3 subpopulations, resulting in reduced proportions of other subpopulations (Figure S5F-G). *Gata2*^R396Q/+^ cells displayed similar behaviors with slight discrepancy characterized by a lesser expansion of MPP2 and MPP3. The kinetic response of mutated HSPCs after LPS stimulation (16 *vs* 96 hours) showed similar patterns across all populations, suggesting that Gata2^R396Q^ mutation might affect the initiation rather than the amplitude of the response (Figure 4H and S5H). Therefore, the *Gata2*^R396Q^ mutation leads to a global hematopoietic functional defect including hyporesponsivess and exhaustion after short- and long-term inflammatory stresses.

### Molecular signatures enriched in *Gata2*^R396Q/+^ LSK cells predict the functional defects

To delve into the molecular mechanisms behind the *Gata2*^R396Q^ mutation-induced functional defects, we adopted an integrative strategy combining chromatin accessibility and gene expression profil analyses on purified LSK cells from 2-month-old *Gata2*^R396Q/+^ and *Gata2*^+/+^ mice (Figure 5A). Over representation analysis (ORA) revealed that the same cellular processes undergo modifications both at chromatin accessibility and expression levels. Mutated LSK cells showed enriched pathways related to migration, inflammatory response, and extracellular matrix degradation, supporting the notion of an exhaustion process (Figure 5B-C and S6A-B). Moreover, the decreased expression and activity of Meis transcription factors, known for their involvement in cell cycle and stem cell self-renewal, in *Gata2*^R396Q/+^ HSPCs might, to some extent, offer a molecular explanation for the exhaustion phenotype (Figure 5D and S6C). The decreased in enrichment of Interferon signaling and erythroblast differentiation pathways (Figure S6A) observed in *Gata2*^R396Q/+^ HSPCs indicates a loss of commitment capacity at steady-state, possibly linked to the reduced binding and expression of Irfs, which are known activators in HSPCs^45, 46^. In *Gata2*^R396Q/+^ LSK, while GATA2 motifs showed only a discreet reduction in their opening, motifs associated with TFs involved in the response commitment, such as Meis^47^ and Irf^48^, displayed significant reductions (Figure 5E). On the other hand, the motifs related to transcription factors involved in cell quiescence (e.g., Egr1) or BMP signaling (e.g., Smad) exhibited the greatest increase in accessibility (Figure 5F). In summary, our findings reveal the transcriptional landscape and network of mutated LSK cells at steady state, which directly correlates with their functional defects observed under challenging conditions. These observations strongly suggest that mutated cells are inherently predisposed to inadequately respond to stimuli.

**Figure 5.**
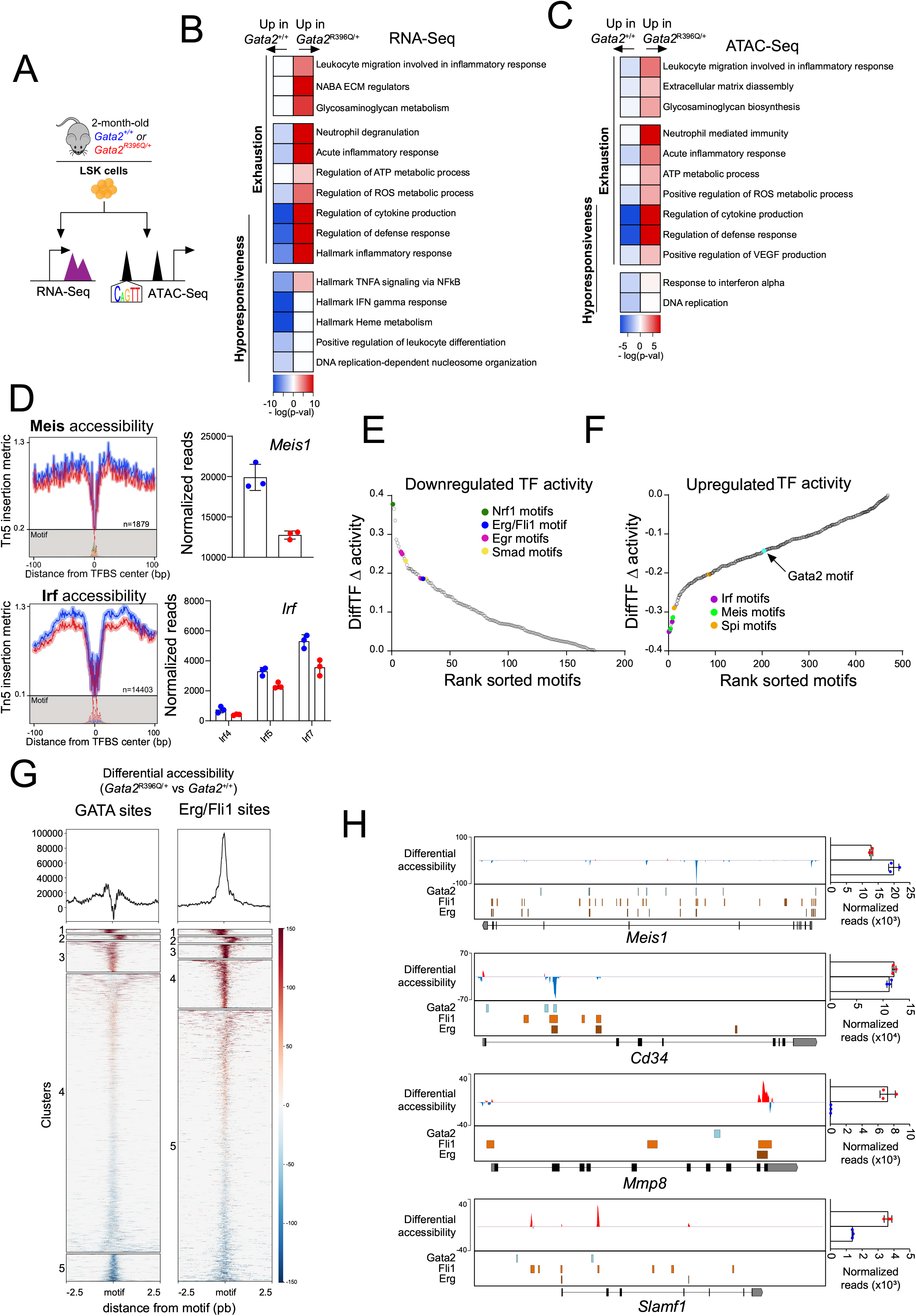
Mutated LSK cells are less prone to respond to stimuli at molecular level. (A) Experimental workflow schematic. (B-C) Heatmap of over-represented analysis (ORA) biological processes in *Gata2*^R396Q/+^ *versus Gata2*^+/+^ cells based on RNA (B) or ATAC (C) sequencing analysis. The color scale represents the enrichment or depletion of each process (-log(p-value)). (D) Digital footprinting analysis at enriched motifs and the number of motifs detected as the frequency of motifs relative to the peak center (values correspond to normalized Tn5 insertions). Meis (top left panels) and Irf (bottom left panels) motifs are shown. The right panels indicate the expression levels of these genes in *Gata2*^+/+^ (blue) or *Gata2*^R396Q/+^ (red) cells. (E-F) Ranked sorted motifs based on differential accessibility: Repressed motifs in *Gata2*^R396Q/+^ LSK cells (E) and enriched motifs (F). (G) ATAC sequencing signal at the Gata or Erg/Fli1 sites. The color scale represents differential accessibility between *Gata2*^R396Q/+^ and *Gata2*^+/+^ conditions. The signals are clustered based on their type: highly more accessible in *Gata2*^R396Q/+^, less accessible, or a mix of the two. The top panel shows the footprint of average accessibility at these sites. (H) The lowest lane represents the genome track indicating exons, 3’ and 5’ UTRs (in grey) of the *Meis*, *CD34*, *Mmp8*, and *Slamf1* (CD150) genes. The top panel indicates higher (red) or lower (blue) accessibility at different loci. The middle panel shows the position of Gata2, Fli1, and Erg binding sites according to ReMap2022^65^ and the expression levels are indicated as histograms at the right of each profile. Each dot represents a pooled sample (red for *Gata2*^R396Q/+^ and blue for *Gata2*^+/+^). Results are shown as mean ± SD; ns: not significant, *p<0.05, **p<0.01, ***p<0.001, ****p<0.0001.

To assess the primary impact of Gata2^R396Q^, our analysis was directed towards the prominent heptad of transcription factors, Scl, Lyl1, Lmo2, Gata2, Runx1, Erg, and Fli1, which collectively define HSPC identity^49^. The discrimination and clustering of regions containing Gata motifs revealed both enrichments and impoverishment of Gata motifs across the genome in mutated LSK cells, explaining why they did not exhibit a pronounced global repression (Figure 5E, G). Moreover, among the heptad motifs, only Erg and Fli1 (ETS motifs) showed a significant change, corresponding to an increase in their activity (Figure 5F-G) in absence of increase expression of Erg, Fli1 or Gata2 (Figure S6F) suggesting a disruption in heptad complex, potentially redirecting these actors to other genomic regions. Accessibility related to ETS-motifs was more specifically located at the promoter than GATA2-motifs underlining a prominent action at the transcriptional level (Figure S6D). To strengthen our hypothesis, we investigated the accessibility to regulatory regions (enhancers and promoters) of these specific motifs located near the loci of genes involved in the hematopoietic process. Regions containing both Gata2 and ETS motifs appeared to be less accessible (Figure 5H and S6G). On the other hand, enhancer and promoter more specifically associated with ETS motif were often more accessible and associated with an increase in expression level of genes such as *Slamf1* (encodes for the CD150) but also *Mmp8* and *Smad1* (Figure 5H and S6E, G).

Our data reinforce the critical role of Gata2 in normal hematopoietic stem cell function at the molecular level, and strongly highlight that its dysfunction affects the ability of hematopoietic stem cells to respond to various stimuli. Finally, our results show some changes in the activities of specific heptad players such as Fli1 and Erg, which have a novel activity in Gata2 mutated context.

### Distinct imbalance of Gata2 allelic expression in the *Gata2*^R396Q/+^ HSPC compartment

We then used flow cytometry to examine the expression level of the Gata2 protein in different subsets of hematopoietic stem and progenitor cells. In *Gata2*^+/+^ mice, we found that Gata2 expression was more prominent in LT-HSCs and MPP2 subpopulations, which aligns with its roles in facilitating self-renewal and influencing erythroid/megakaryocyte differentiation bias in these subpopulations, respectively. In *Gata2*^R396Q/+^ mice, Gata2 protein expression was predominantly higher in LT-HSCs, MPP2, and MPP3 subpopulations compared with *Gata2*^+/+^ mice (Figure 6A and S7A). We then explored the impact at the protein level of one of Gata2’s canonical target genes, Gata1. Since Gata1 protein expression is barely detectable in LSK, we focused our investigation on the LK compartment and noted reduced Gata1 protein levels in MEP cells of *Gata2*^R396Q/+^ mice (Figure S7B) suggesting that *Gata2* R396Q mutation may impact the expression of Gata2 target genes (Figure 6B and S7C). The variations in GATA2 protein expression across different hematopoietic stem and progenitor cell (HSPC) subpopulations may help to understanding the diverse range of defects observed within each of these subsets.

**Figure 6.**
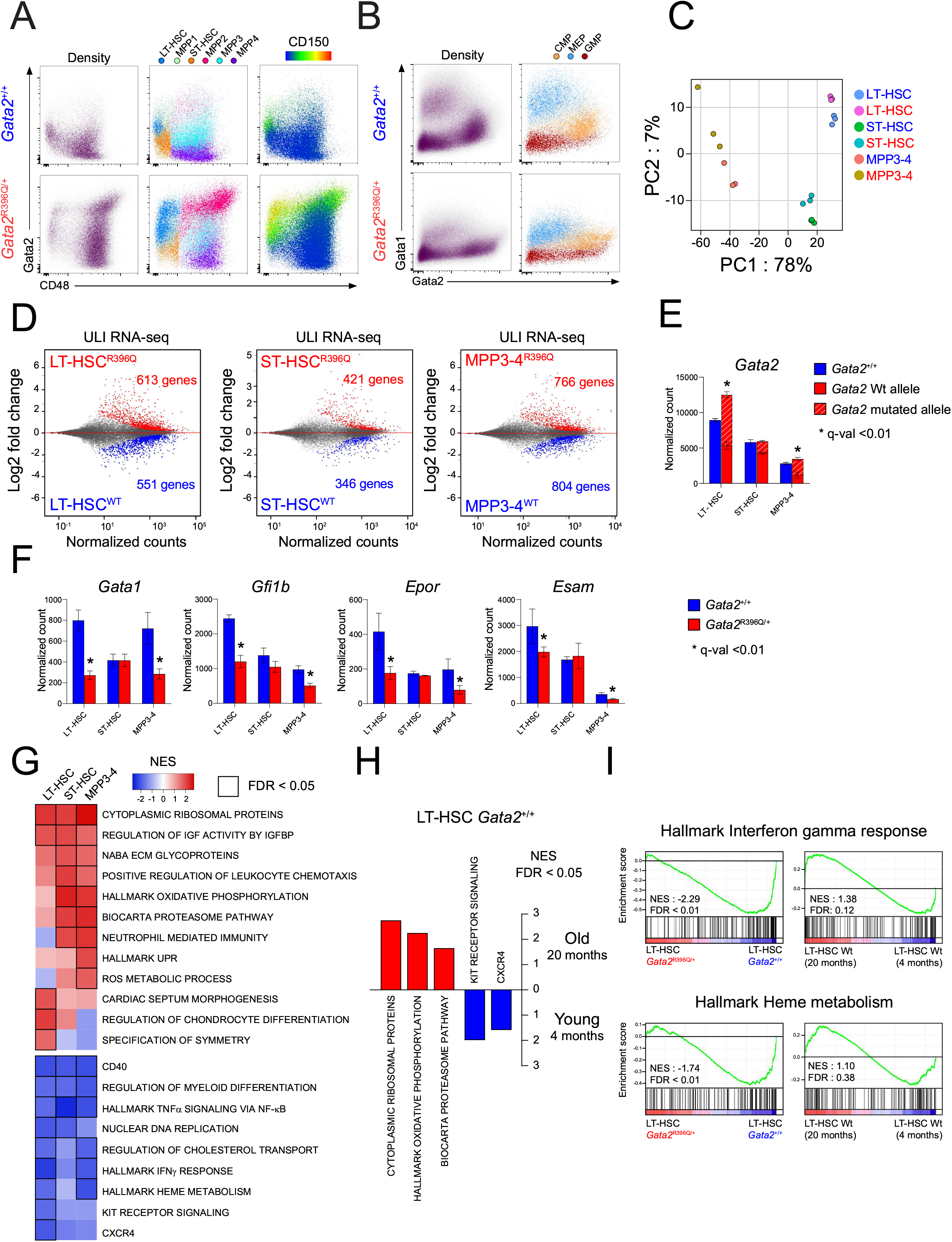
Differential expression of the mutated allele induces specific signature in LSK subpopulations. (A) TriMap representing intracellular FACS analyses of Gata2 protein on LSK subpopulations of *Gata2^+/+^* and *Gata2*^R396Q/+^ bone marrow (n=3). CD48 marker is shown on the x-axis and CD150 marker intensity is color-coded. (B) Dot plots showing intracellular Gata1 and Gata2 protein expression in subpopulations of *Gata2^+/+^* and *Gata2*^R396Q/+^ LK cells (n=3). (C) Principal component analysis of *Gata2*^+/+^ (blue) and *Gata2*^R396Q/+^ (red) LT-, ST-HSC and MPP3-4 cells. (D-E) MA-plots illustrating fold difference relative to normalized read counts for RNA-sequencing reads, depicting significantly increased (red), significantly decreased (blue), or unchanged (grey) expression in *Gata2*^R396Q/+^ *versus Gata2*^+/+^ LT-HSC (D, left panel), ST-HSC (D, middle panel), and MPP3-4 (D, right panel) subpopulations of LSK cells. (E) Expression of *Gata2* in *Gata2*^+/+^ (blue) or *Gata2*^R396Q/+^ (red) LT-ST-HCS and MPP3-4 cells (F) Expression of *Gata1, Gfi1b, EpoR* and *Esam* genes in *Gata2^+/+^* (blue) or *Gata2*^R396Q/+^ (red) LT-, ST-HCS and MPP3 cells. Results are shown as mean ± SD; *q-val<0.01. (G) Gene set enrichment analysis (GSEA) of enriched or depleted Gene Ontology (GO) terms for LT-, ST-HSC, and MPP3-4 cells, represented by a color scale of normalized enrichment score (NES). False discovery rates (FDR) <0.05 are indicated by bold-framed squares. (H) Hallmark-term analysis of gene sets enriched in young *Gata2^+/+^*LT-HSCs *versus* 20-month-old *Gata2^+/+^* LT-HSC cells, with normalized enrichment scores (NES) shown for significant pathways (FDR <0.05). (I) GSEA enrichment plots for interferon response (top panel) and heme metabolism (bottom panel) comparing *Gata2*^R396Q/+^ to *Gata2^+/+^* LT-HSC (left panels) and old LT-HSC *versus* young LT-HSC (right panels).

To dig further into the molecular consequences of the mutation, RNA-Seq analysis was performed on sorted LT-HSC, ST-HSC, and MPP3-4 subpopulations. Principal component analysis (PCA) clustering illustrated that the mutant expression did not significantly alter the molecular identity of each population, as they remained closely clustered with their wild-type counterparts (Figure 6C). Nonetheless, the number of differentially expressed genes between mutated and non-mutated cells was higher in LT-HSCs and MPP3-4 than in ST-HSCs (Figure 6D). This increase seemed to be proportionally associated with the ratio of the mutated Gata2 allele, higher in LT-HSCs and MPP3-4 compared to ST-HSCs (58%, 67% and 28% respectively, Figure 6E) highlighting an allele-specific expression (ASE). Furthermore, the expression of established Gata2 target genes exhibited variability based on the expression level of the mutated allele. Remarkably prominent canonical Gata2-activated genes, such as *Gata1* and *Gfi1b*, demonstrated significant expression reductions exclusively in LT-HSCs and MPP3-4, while Gata2-repressed *Cebpa* gene^50^, displayed an increase within these populations. Moreover, other genes critical for HSPC functions, such as *Epor*, *Esam*, *Aqp9*, and *Gpc3*, exhibited specific downregulation in LT-HSCs and MPP3-4 (Figure 6F and S7D-E).

To deepen our understanding of the altered transcriptomic landscape within each population, we conducted clustering analysis of GSEA scores using KEGG, GO BP, and Hallmark pathway databases. The results of this analysis unveiled notable dysregulation of pathways related to protein synthesis, myeloid differentiation, and TNFα response in sorted subpopulations to a comparable degree, regardless of the ASE status. This suggests that these pathways undergo significant perturbations independent of the allelic expression patterns. Conversely, repression of the IFN gamma response and heme metabolism pathways notable disruption solely in LT-HSCs and MPP3-4 exhibiting higher expression of mutant allele than wild-type allele. Additionally, certain dysregulated pathways appeared to be more confined to mutated LT-HSCs encompassing pathways related to symmetry specification, Kit receptor signaling and even pathways unrelated to hematopoiesis (Figure 6G).

Given the cellular phenotypic characteristics reminiscent of a premature aging phenotype, we conducted an investigation to ascertain whether the modifications induced by the mutation in *Gata2*^R396Q/+^ mice might derive form molecular aging-related features similar to the observed phenotypic increase of LT-HSC with high CD150 expression level^51^ at a younger age. We therefore reanalyzed RNA-Seq data from Maryanovich and colleagues^52^ from aging cells using the same pipeline employed in our study. We highlighted in 20-month-old wild-type LT-HSCs and *Gata2*^R396Q/+^ HSPCs similar pathways deregulations, including enrichment for oxidative phosphorylation and proteasome-related pathways, as well as a loss of enrichment for signaling pathways associated with Kit and CXCR4 receptors (Figure 6H). Notably, interferon signaling and inflammatory response are reported to be involved in aging process^48^. Interestingly, we observed a significant increase in the expression of specific genes from these pathways in 20-month-old LT-HSCs (Figure 6I), indicating the potential involvement of Gata2 in these pathways during the aging process.

Our findings emphasize that the impact of relative amount of mutant varies significantly across different molecular pathways. Notably the increased expression of the mutated allele exerts a significant influence on essential pathways related to stimulus responses and HSPC differentiation. These observations underscored the intricate regulatory interplay between the mutant allele and crucial molecular processes, offering valuable insights into the underlying mechanisms governing HSPC function and contributing to our understanding of hematopoietic disorders.

### Allele-Specific Expression mechanism drives functional defects in LT-HSC subpopulation

Impairment of LT-HSCs, the cells at the top of the hematopoietic hierarchy, disrupts hematopoietic homeostasis. To elucidate the transcriptomic deregulations in this population, given the observed functional defects in mutated LT-HSCs, we used molecular signatures associated with hallmark HSCs properties. The MoIO^53^ signature, linked with superior HSC function, is overrepresented in unmutated LT-HSCs as well as the NoMO^53^ signature which is associated with less quiescent and functionally inferior HSCs (Figure 7A). Notably, specific genes within the MolO signature like *Neo1*, *Smtnl1*, *Cd82* and *Vwf* exhibited reduced expression in mutated LT-HSCs whereas *Ly6a* and *Procr* genes encoding Sca-1 or Epcr respectively showed increased expression (Figure 7B and S7F). Similarly, most of the NoMO signature genes, including *Muc13*, *Itga2b*, *Gata1* or *Aqp9*, displayed decreased expression in mutated LT-HSCs (Figure 7B and S7G). Additionally, the expression of *Dlk1*, a gene associated with heightened myeloid potential, quiescence state, and self-renewal capacity^54^, was notably decreased in mutated LT-HSCs (Figure 7B). Furthermore, mutated LT-HSCs expressed genes typically not found expressed in hematopoietic cells such as *Igf1*, *Sfrp2*, *S1pr3 or Nsg2* (Figure 7C and S7H). These findings indicated that mutated LT-HSCs undergo a loss of general HSC features, encompassing both dormant and active HSC properties.

**Figure 7.**
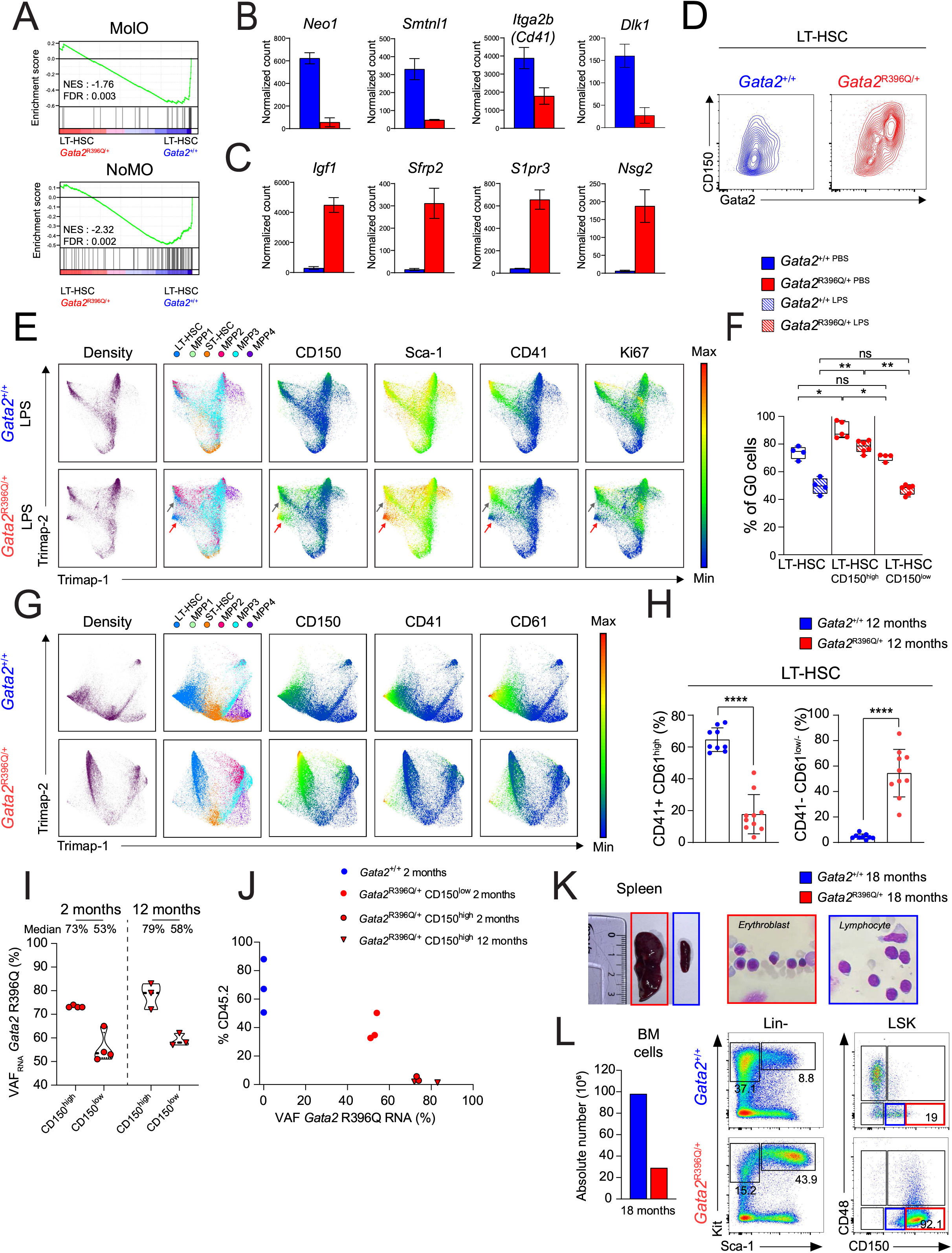
LT-HSC population is heterogeneous and associated with the existence of aberrant cells. (A) GSEA-enrichment plots showed significant depletion of Molo and NoMO signatures^53^ in LT-HSC form *Gata2^R396Q^*^/+^ mice. (B) Expression of *Neo1, Smtnl1, Itga2b* and *Dlk1* genes (from the Molo and NoMO signature), in *Gata2*^+/+^ (blue) or *Gata2*^R396Q/+^ (red) LT-HSCs. (C) Expression of non-hematopoietic related genes. (D) Flow cytometry analysis of Gata2 protein expression according to CD150 expression in LT-HSCs from 2-month-old *Gata2*^+/+^ (blue) and *Gata2*^R396Q/+^ mice. (E) TriMap visualization of bone marrow cells from PBS- or LPS-injected 2-month-old *Gata2*^+/+^ (bleu) and *Gata2*^R396Q/+^ (red) mice depicting HSPC subpopulations, population density and CD150, Sca-1, CD41 and Ki67 marker intensity of on each population. Intensity of each marker is shown using a colored scale. (F) Percentage of cells in G0 (Ki67^-^ cells) of total LT-HSC *in Gata2*^+/+^ (blue) or CD150^high^ and CD150^low^ LT-HSCs in *Gata2*^R396Q/+^ (red) mice injected with PBS (plain box) or with LPS (hatched box). (G) TriMap visualization on the different LSK subpopulations showing the population density and the intensity of the CD150, CD41 and CD61 markers on each population following a colored scale. (H) Statistics of absolute number of CD41/CD61 double positive (CD41^+^/CD61^+^) or CD41^-^/CD61^-/low^ LT-HSCs. (I) Variant allelic frequency at RNA level (VAF_RNA_) of *Gata2* R396Q mutation on sorted CD150^low^ and CD150^high^ LT-HSCs from 2- and 12-month-old mice. (J) Analysis of CD45.2 cells percentage after 1 month. 500 CD45.2 LT-HSCs from *Gata2*^+/+^ mice (2-month-old mice n = 3, blue), *Gata2*^R396Q/+^ mice (2-month-old: red circles (CD150^high^: black box), 12-month-old: red triangle (CD150^high^: black box)) were transplanted in CD45.1 recipient mice. (K) Splenic extramedullary hematopoiesis of 18-month-old-*Gata2*^R396Q/+^ mouse (red) compared to *Gata2*^+/+^ mouse (blue) at the same age. Spleen size (left panel), cytospin of collected splenic cells stained with May-Grünwald-Giemsa coloration (right panel). (L) Number of bone marrow cells (left panel) and visualization of LSK subpopulations by flow cytometry from 18-month-old-mice (*Gata2*^+/+^ in blue, *Gata2*^R396Q/+^ in red). Results are shown as mean ± SD; *p<0.05, **p<0.01, ***p<0.001, ****p<0.0001.

The LT-HSCs population is nowadays recognized heterogeneous^20^. In *Gata2*^R396Q/+^ LT-HSCs, we observed two phenotypically distinct populations based on Gata2 and CD150 protein levels (Figure 7D). To explore *Gata2*^R396Q/+^ LT-HSCs functional heterogeneity, we subjected the mice to acute inflammatory stress by LPS. This assay revealed two distinct LT-HSCs subpopulations in *Gata2*^R396Q/+^ mice (red and grey arrows, Figure 7E). Sixteen hours after administering LPS, the majority of *Gata2*^+/+^ LT-HSCs expressed the platelet marker CD41 (encoded by *Itga2b*), while only a fraction of *Gata2*^R396Q/+^ LT-HSCs exhibited its expression (grey arrow). Moreover, a distinct aberrant LT-HSC subpopulation exclusively found in *Gata2*^R396Q/+^ mice displayed a higher CD150 and Sca-1 markers intensities and a lower of Kit intensity (red arrow, Figure 7E and S8A-B). This phenotypic distinction between these two types of LT-HSCs in *Gata2*^R396Q/+^ mice enabled us to reveal that CD150^high^ aberrant LT-HSCs cycled less than CD150^low^ and WT LT-HSCs at steady state. Besides, in contrast to CD150^low^ and WT LT-HSCs, which showed a cycling phenotype under the effect of LPS, (Figure 7F), Ki67^-^ cell proportion remained rather unchanged in the CD150^high^ LT-HSCs population.

Physiological aging induces phenotypic changes in LT-HSCs such as an increased expression of CD150 marker and detection of megakaryocytic surface markers such as CD41 or CD61^42, 55^. This platelet bias in LT-HSCs correlates with the upregulation of *Gata2* (Figure S8C) and its target gene *Gata1*^56^. Leveraging a combination of CD150, CD41 and CD61, we were able to discriminate the aberrant CD150^high^/CD41^-^/CD61^-/low^ LT-HSCs in middle-aged *Gata2*^R396Q/+^ mice, which expanded over time and represented the majority of the LT-HSCs population. (Figure 7G-H and S8D). These distinct LT-HSCs exhibited hypofunctionality due to their loss of clonogenic capacity and multipotency as evidenced by a significant reduction in CFU-GEMM and CFU-Mk colony number, which correlated with the absence of CD41 protein expression (Figure S8E-F).

Moreover our aim was to further apprehend the existence of these 2 populations in *Gata2*^R396Q/+^ mice. Building upon our previous discovery of an ASE mechanism in HSPC subpopulations, we delved into the prospect of different allelic imbalances in these two subpopulations. To explore this, we sorted CD150^high^ and CD150^low^ LT-HSCs from 2- and 12-month-old *Gata2*^R396Q/+^ mice and measured their variant allelic frequency (VAF) of the Gata2 mutation at the RNA level. Our results revealed that *Gata2*^R396Q^ VAF_RNA_ was higher in CD150^high^ compared to CD150^low^ LT-HSCs (73% *vs* 53% at 2 months respectively) providing evidence for an ASE mechanism in the CD150^high^ LT-HSCs (Figure 7I).

In an additional LT-HSC subpopulation transplantation assay, we observed an inverse relationship between the level of donor chimerism and *Gata2* VAF_RNA_. Transplantation of CD150^high^ LT-HSC subpopulation resulted in a severe repopulation defect in recipient bone marrow one-month post-injection (Figure 7J), confirming the hypofunctional nature of the CD150^high^ LT-HSCs. On the other hand, transplantation of CD150^low^ underscored a milder defect in BM repopulation capacity compared to *Gata2*^+/+^ LT-HSCs.

Interestingly, these functional defects were detected in old mice at steady-state. Indeed, the phenotypic characterization of an 18-month-old *Gata2*^R396Q/+^ mouse revealed a significantly reduced hemoglobin level (6g/dL compared with 10g/dL for the *Gata2*^+/+^ mouse of the same age), an enlarged spleen size with extramedullary erythropoiesis and a substantial decreased in the number of bone marrow cells (Figure 7K-L). Intriguingly, the LSK compartment was predominantly composed of CD150^high^ LT-HSC cells suggesting that bone marrow hypoplasia could be attributed to the exhaustion of the most functional LT-HSCs (CD150^low^).

Additionally, we reported the case of a patient bearing the germline missense *GATA2* R362G (c.1084C>G) mutation, diagnosed with AML. Our RNA-Seq data confirmed ASE with a higher relative expression of the mutated allele (Figure S8G). These findings underscore the clinical relevance of the ASE mechanism in human GATA2 deficiency syndrome and its potential implications for disease development and progression.

In summary, our results demonstrate that *Gata2*^R396Q^ mutation leads to a loss of HSCs’ stemness identity, resulting in functional defects. Notably, we observed accumulation of an aberrant population characterized by a mutated allele-specific expression (ASE) mechanism. This finding closely mirrors the ASE phenomenon observed in GATA2 deficient patients at distinct stages of disease progression^22^. Thus, our study provides a valuable model that recapitulates the ASE-related pathology observed in *Gata2* deficiency, shedding light on the underlying mechanisms contributing to disease development and transformation.

## Discussion

This study offers evidence elucidating the effects of the *GATA2*^R396Q^ germline mutation on the stem cell compartment, using a *Gata2*^R396Q/+^ knock-in mouse as an experimental model. Notably, when compare with the *Gata2*^+/-^ or Gata2 +9.5^+/-^ mouse models^57^ both exhibiting lower expression of Gata2, the *Gata2*^R396Q/+^ and *Gata2*^R398W/+^ murine models with missense mutations demonstrate no reduction in overall Gata2 expression in the LSK compartment^17^. This observation suggests the existence of mechanistic variations depending on the specific type of mutation. Furthermore, we have demonstrated that the missense GATA2^R396Q^ mutant retains its nuclear localization, in contrast to a truncated variant. This preservation of its nuclear localization may suggests that the GATA2^R396Q^ variant might play an aberrant role, potentially affecting the hematopoietic program^15, 58^. Despite the observed decrease of its DNA binding activity^37, 58^ our findings suggests that GATA2^R396Q^ may still hold functional implications within this cellular context. However, the lack of viable *Gata2*^R396Q/-^ mice indicates that the Gata2^R396Q^ variant are unable to rescue the phenotype due to the loss of the wild-type allele, and it functions as a global loss-of-Gata2 function variant. Nonetheless, we have demonstrated that *Gata2*^R396Q/-^ embryos, like *Gata2*^R398W/R398W^ embryos present delayed lethality compared with *Gata2*^-/-^ embryos^3^ supporting the notion that Gata2^R396Q^ is not a null-mutant.

Molecular analyses of HSPCs subpopulations from *Gata2*^R396Q/+^ mice revealed a noticeable reduction in the expression of Gata2 target genes. However, the overall Gata2 activity did not show significant variation, as the number of more accessible was equivalent to the number of less accessible sites Furthermore, despite the reduction of the 9.5kb enhancer region accessibility, we did not observe a reduction in Gata2 expression in bulk LSK. These findings raise some questions about the underlying mechanisms governing the fine regulation of Gata2 expression and activity.

Gata2 forms a heptameric complex with six other transcription factors, including Runx1, Lyl1, Erg, Fli1, Lmo2 and Tal1^49^. Our observations indicate that the Gata2^R396Q^ mutant disrupts DNA binding specifically at the heptamer binding site, while leaving the expression of other complex’s transcription factor unaffected. In particular, we observed a significant increase in the accessibility only at Fli1/Erg motifs among heptamer transcription factors. The more accessible binding sites of these factors are located close to gene promoters leading to enhanced expression of specific genes, such as the *Smad* genes involved in the BMP pathway. Noteworthy, Smad1 and Smad5 have been implicated in Gata2 expression^59, 60^. This suggests a potential interplay between the Smad signaling pathway and Gata2, which may contribute to maintain overall Gata2 expression at levels close to those observed in wild-type mice. Similarly, the *Slamf1* gene, which encodes the CD150 marker, has been shown to be correlated with Gata2 expression in normal hematopoietic stem cells (HSCs) through Akt, p38 MAPK, or Erk pathways^50^. Interestingly, we observed an increased accessibility of Fli1 sites in mutated HSPCs at *Slamf1* loci suggesting possible adaptive response initiated by the cells to compensate for the reduced Gata2 activity resulting from the mutation. These findings highlight the intricate interplay between Gata2, BMP signaling, and CD150-mediated pathways, emphasizing the complex regulatory mechanisms governing HSPCs and their ability to adapt in response to genetic perturbations.

The *Gata2*^R396Q/+^ mice displayed an abnormal distribution of LSK subpopulations, a characteristic that was evident as early as embryonic stages and remained present throughout their lifespan. Notably, at steady-state conditions, the *Gata2*^R396Q/+^ mice unveil an increased proportion of LT-HSCs, which is an unique feature of this model in comparison to the *Gata2*^+/-^ and *Gata2*^R398W/+^ models, which show a reduction or no significant difference in proportion respectively. Nonetheless, all models display a reduction in ST-HSCs^5, 17, 61^. Despite an increase in the number of LT-HSCs, our findings provide compelling evidence that the molecular consequences of the mutation lead to impaired functionality of hematopoietic stem and progenitor cells (HSPCs) under challenging conditions. We observed significant enrichments of inflammatory and mobilization-related molecular pathways that appeared to counteract self-renewal mechanisms, as evidenced by the downregulation of Meis1 expression and activity in the mutated HPSCs. These findings shed light on the underlying reasons for the reduced engraftment and long-term persistence capabilities of mutated cells to engraft and persist over a long period. Furthermore, the LPS stimulation assay demonstrated a widespread hyporesponsiveness across all subpopulations with the exception of ST-HSCs. This unique characteristic of the ST-HSCs prompted us to investigate the overall abundance of Gata2 protein within all LSK subpopulations. Interestingly, we observed a significant increase in global Gata2 protein specifically in the mutated subpopulations characterized by higher cell numbers (LT-HSCs, MPP2 and MPP3). In contrast, minimal variation in Gata2 protein expression was observed in the less affected populations, namely the ST-HSCs. Taken together, these experimental findings strongly suggest a correlation between the disturbed equilibrium of HSPCs subpopulations and the expression level of Gata2.

The MolO molecular signature associated with the dormant HSC phenotype was found to be enriched in non-mutated LT-HSCs. In contrast, the mutated LT-HSCs showed a significant decrease in the expression of *Neo1* (as part of the MolO signature). Additionally, the NoMO signature, which is associated with less quiescent, functionally inferior HSCs was enriched in *Neo1*-mutant HSC with an increase of Gata1 and Itga2b expressions^62^. In our *Gata2*^R396Q/+^ mouse model, *Gata1*, being a target gene of Gata2, was unable to undergo its physiological upregulation, resulting in the absence of enrichment of the NoMO signature in mutated LT-HSCs. Furthermore, the mutated LT-HSCs expressed genes that are typically expressed in microenvironment cells rather than in hematopoietic cells. This observation is consistent with the findings from the De Pater’s group^63^, which demonstrated impairment in suppressing endothelial identity in *Gata2*^+/-^ HSPCs during embryonic maturation due to a reduced Gfi1b activity. This reduction was also observed in *Gata2*^R396Q/+^ LT-HSCs. Collectively, our findings reveal a loss of molecular stemness identity of mutated HSCs with alterations in gene expression patterns that may impact their quiescence and functional properties.

In *Gata2*^R396Q/+^ mice, we noticed a higher presence of CD150^high^-expressing LT-HSCs cells compared to wild-type mice, showing similarities to certain aspects of an aging phenotype. Both aged LT-HSCs and mutated HSPCs displayed dysregulation in pathways related to protein synthesis and oxidative phosphorylation. However, there were notable differences in the gene expression profiles of LT-HSCs and MPP3-4, which express the mutated allele in higher proportions. These specific subpopulations showed a global downregulation of the genes associated with interferon and heme metabolism pathways, contrasting with the context of aging where certain of these genes are upregulated. These findings suggest that Gata2 plays a role in physiological aging and indicates potential distinct mechanisms underlying the effects of the Gata2^R396Q^ mutation and aging on HSCs and HSPCs^52^.

Additionally, in our study we have uncovered significant heterogeneity within the HSC population, characterized by the presence of phenotypically and functionally aberrant LT-HSCs. These cells show an allelic bias, favoring the expression of the mutated allele. This ASE mechanism has previously reported in the context of GATA2, being more frequent in comparison to other genes associated with myeloid or cancer-related conditions^21^. Interestingly, Al Seraihi and colleagues have reported that differential allele expression can influence the clinical phenotypes in a GATA2 deficient patient^22^. Our findings showed that ASE is present in mice as early as 2 months old and might persist even in the context of a transformed phenotype in the patient. This raises questions about potential secondary events that could favor the selection of cells demonstrating ASE. Furthermore despite the existence of genotype-phenotype correlations^64^, there remain questions regarding the wide range of symptoms observed even within the same family. The investigation of ASE mechanism could provide valuable insights into understanding these divergences and shed light on the underlying factors contributing to the variable clinical manifestations associated with *GATA2* mutations.

In conclusion, our study provides compelling evidence for the substantial impact of a missense *Gata2* mutation on the HSPC compartment during hematopoietic development. The alterations at phenotypic, functional, and molecular levels highlight the crucial role of Gata2 in maintaining HSPC homeostasis. The disruption of critical molecular pathways in mutated HSPCs hampers their ability to respond appropriately to environmental signals and may contribute to their exhaustion. These insights significantly advance our understanding of hematopoiesis and lay the groundwork for future investigations into the pathogenesis and therapeutic approaches for disorders associated with *GATA2* mutations. Therefore, our findings hold promising implications for improving clinical follow-up for patients with GATA2-related conditions, as they provide valuable insights into the mechanisms involved in maintaining HSPC homeostasis.

## Acknowledgments

We greatly thank Prof. Stuart Orkin for the *Gata2*^+/-^ mice and Prof. Emery Bresnick for the Gata2 antibody. We are grateful to Manon Farcé from the cytometry and cell-sorting facility of the Pole Technologique of the CRCT (INSERM U1037) for technical assistance. We thank the Anexplo/Genotoul platforms for technical assistance (UMS006). L.J. was supported by la Société Française des Cancers de l’Enfant (SFCE, association Capucine, R17094BB) and Région Occitanie grants (RPH17006BBA). This specific project was founded mainly by INCa (R17164BB and R21196BB), association Laurette Fugain (R20110BB), Fédération leucémie espoir (R2004BB), and Association « Constance la petite guerrière astronaute » (R19043BB) and association « les 111 des arts » (R18062BB). The team is « Equipe labélisée Ligue Contre le Cancer » and was also supported by la Ligue Nationale Contre le Cancer (R19015BB), Société Française des Cancers de l’Enfant (R17094BB, association Capucine), la Région Occitanie (R16038BB), Association Cassandra (R18041BB).

## Author contributions

L.L., L.J., V.F., C.H., E.S., P.E., M.B., S.H. and C.D. performed and analyzed *in vivo* and *in vitro* studies. N.P. and S.D. performed library preparation. L.L., L.J. and V.F analyzed the data from RNA-Seq and ATAC-Seq. L.L., V.F., B.G., C.D., E.D. and C.B. conceived and designed the experiments. E.D., M.P and C.B. acquire the funding. L.L., L.J., V.F. and C.B. wrote the manuscript with feedback from all authors.

## Disclosure of Conflicts of Interest

The authors declare no competing financial interests.

## Notes

### Competing Interest Statement

The authors have declared no competing interest.

