## Supplemental material for "Loss of HSC stemness identity is associated with exhaustion and hyporesponsiveness in GATA2 deficiency syndrome"

#### Figure S1

(A) Immunofluorescence on Ba/F3 cells transduced either with a GFP-expressing vector (named empty vector) or with GFP and wild type GATA2 (GATA2 WT) or with two Gata2 mutants: GATA2 R396Q and GATA2 R204X. GATA2 proteins had been tagged with a HA tag. Are shown representative pictures of the GFP fluorescence (Top Lane), of the Dapi labeling (second lane) of the anti- HA-RFP fluorescence (third lane) and the merge of Dapi and HA-RFP fluorescence (fourth lane), the scale is indicated in the first left picture, top lane. At the bottom of each column are representative histograms of the Dapi and RFP fluorescence along a section of the cell (drawn in white in the merge pictures) indicating the co-localization of the signals. (B) Ratio of the cytoplasmic *versus* nuclear RFP fluorescence showing that GATA2 WT and GATA2 R396Q are exclusively nuclear as GATA2 R204X are mainly cytoplasmic (n>10 cells; Zen software (Zeiss). Results are shown as mean ± SEM. Unpaired t-test. *p<0.05, **p<0.01. (C) Schematic diagram of the protocol for studying the ectopic expression of GFP (MIG) or wild type GATA2 (GATA2 WT) or mutant GATA2 (GATA2 R396Q) in murine lineage negative ($Lin^-$) cells. The different steps and analyses are indicated on a timeline. (D) Gating strategy for identifying the cells remaining after methylcellulose cultures. (E) Scheme of the steps to obtain the knock-in *Gata2*$^{R396Q/+}$ mouse model. The first line shows the genomic organization of the *Gata2* gene. X indicate the exons. The second line depicts the homologous recombination vector. The third line shows the resulting allele after the homologous recombination. The fourth line shows how the Neo resistant gene, flanked by two FRT sites (bleu arrows), has been removed with Flp deletion in Flp deletor mice. The fifth line show the result of the CRE recombination and the germline transmission of this knock-in allele to obtain a constitutive knock-in mouse model after breeding with Vav-Cre mice. 2 sets of different LoxP sites (wt and 577) and their orientations are indicated by colored triangles. (F) Representative examples of embryo (E) genotyping detecting by PCR a 592pb wild-type allele, a 796pb of *Gata2* R396Q allele (top panel) and a band from the Neomycin Resistant *(Neo)* gene from the recombinant knock-out allele (bottom panel). On these examples only the E5 and E6 bare the *Gata2*$^{R396Q}$ allele and E2, E4 and E6 are on a Gata2 deficient background. (G) Pictures of a *Gata2*$^{R396Q/-}$ embryo showing a delay in differentiation. (H) Genotype of the 18 embryos from two litters. Y: presence of the allele, N: absence of the allele. The bottom line provides the genotype.

#### Figure S2

(A) Representative FACS analysis of 14.5 days LSK fetal liver cells with CD48, CD34 and CD150 markers *Gata2*$^{+/+}$ (in blue, n=3) and *Gata2*$^{R396Q/+}$ (in red, n=3) embryos. (B) Expression of CD150 and Sca-1 markers on LSK of 2-month-old *Gata2*$^{+/+}$ (in blue) and *Gata2*$^{R396Q/+}$ (in red). (C) Percentage of cells in G0 (Ki67$^{-}$ cells) of each subpopulation of hematopoietic stem and progenitor cells of 2-month-old *Gata2*$^{+/+}$ (in blue) and *Gata2*$^{R396Q/+}$ (in red). (D-E) Gating strategy and representative example of FACS analysis of 2-month-old *Gata2*$^{R396Q/+}$ (D) or *Gata2*$^{+/-}$ (E) bone marrow LSK cells and their wild-type counterparts. (F) Absolute number of the different compartments of hematopoietic and progenitor cells of 2-month-old wild-type *Gata2*$^{+/+}$ (in blue) and *Gata2*$^{+/-}$ (in green) mice. (G) CD150 mean of fluorescence (MFI) in *Gata2*$^{+/+}$ (in blue) and *Gata2*$^{+/-}$ (in green) LT-HSC cells.

Lin$^{-}$: lineage negative cells, LK: Lin$^{-}$Kit$^{+}$, LSK: Lin$^{-}$Sca-1$^{+}$Kit$^{+}$; LT-: long-term- and ST-: short-term- HSC: hematopoietic stem cells; MPP: Multi Potent Progenitor. Each dot represents an individual mouse. Results are shown as mean ± SD; Unpaired t-test. *p<0.05, **p<0.01, ***p<0.001, ****p<0.0001, ns: non-significant.

#### Figure S3

(A) TriMap on Lin$^{-}$ Kit$^{+}$ cells of 2-month-old *Gata2*$^{+/+}$ (top line) and *Gata2*$^{R396Q/+}$ (bottom line) showing the density and the intensity of the CD150, Kit, Sca-1, CD34, CD48 and CD16/32 markers on each population following a colored scale. (B) Gating strategy for LK subpopulations including CMP, GMP and MEP (C) Representative dot plot of CD34 and CD16/32 marker expression at the surface of LK cells of 2-month-old *Gata2*$^{+/+}$ (in blue), *Gata2*$^{R396Q/+}$ (in red) and *Gata2*$^{+/-}$ (in green) mice. (D) Statistics of FACS analyses of the absolute number of LK, CMP, GMP and MEP cells in bone marrow of 2-month-old *Gata2*$^{+/+}$ (in blue, n=15) and *Gata2*$^{+/-}$ (in green, n=16) mice. (E) proportion as pie charts showing the distribution of B (purple), T (green) and myeloid (orange) cells in the bone marrow of 2-month-old *Gata2*$^{+/+}$ (bleu dots) and *Gata2*$^{R396Q/+}$ (red dots) mice. (F-G) Blood parameters of 2-month-old *Gata2*$^{+/+}$ (in blue) and *Gata2*$^{+/-}$ (in green) mice (F) and 12-month-old *Gata2*$^{+/+}$ (in blue) and *Gata2*$^{R396Q/+}$ (in red) mice (G).

LK: Lin$^{-}$Kit$^{+}$, CMP: Common myeloid progenitor, MEP: Megakaryocyte–erythroid progenitor, GMP: Granulocyte-monocyte progenitor; MPP: Multipotent progenitor. Hb: Hemoglobin WBC: white blood cells. Each dot represents an individual mouse. Results are shown as mean ± SD; ns: not significant, *p<0.05, **p<0.01, ***p<0.001, ****p<0.0001.

#### Figure S4

(A) Clonogenic assay on LSK cells of 2-month-old *Gata2*$^{+/+}$ (in blue) *Gata2*$^{R396Q/+}$ (in red) mice. LSK cells were plated on cytokine-enriched methylcellulose medium. Colonies were counted and cells were serially plated every 7 days (n=6) for 3 passages (I, II and III). (B) Representative FACS analysis of cells recovered from methylcellulose after the third cell plating. (C) May Grunwald Giemsa coloration of cytospin of cells recovered from methylcellulose after the third cell plating of *Gata2*$^{+/+}$ cells (upper panel) and *Gata2*$^{R396Q\ /+}$ (lower panel). (D) Clonogenic assay on 2-month-old LSK cells of *Gata2*$^{+/+}$ (in blue) *Gata2*$^{+/-}$ (in green) mice. LSK cells were plated on cytokine-enriched methylcellulose medium. Colonies were counted and cells were serially plated every 7 days (n=3) for 3 passages (I, II and III). (E) Colonies were identified and quantified 10 days after the first cell plating. (F) Representative dot plot of FACS on cells recovered from methylcellulose after the third cell plating (left panel) and May Grunwald Giemsa coloration of cytospin (right panel) of cells recovered from methylcellulose after the third cell plating of *Gata2*$^{+/+}$ cells (upper panel) and *Gata2*$^{+/-}$ (lower panel). CFU: Colony forming unit; GEMM: Granulocyte, erythroid, macrophage, megakaryocyte; GM: Granulocyte, macrophage; G: Granulocytes; M: Macrophage and BFU/CFU-E-MK: Burst-forming unit/ Colony forming unit Erythroid-Megakaryocytes. Results are shown as mean ± SD; ns: not significant, *$p<0.05$, **$p<0.01$, ***$p<0.001$, ****$p<0.0001$.

#### Figure S5

(A) Engraftment quantification of total bone marrow CD45.2$^{+}$ cells (from *Gata2*$^{+/+}$ in bleu of *Gata2*$^{+/-}$ in green as donor mice) recovered 2 or 6 months after the syngeneic transplantation; the number of mice which are considered engrafted (≥1% of CD45.2$^{+}$ cells) out of the total number of transplanted mice and the median of engraftment percentages are indicated on the top of the graph. Right panel indicates the percentage of CD45.2$^{+}$ cells 6 months after a secondary transplant. Each blue dot represents an individual mouse. (B-C) Pie charts showing relative proportions of bone marrow myeloid (orange), B (purple) and T (green) lymphoid cells into the CD45.2$^{+}$ population at 2 and 6 months after engraftment from *Gata2*$^{+/+}$ and *Gata2*$^{+/-}$ total bone marrow (B) or *Gata2*$^{R396Q/+}$ and *Gata2*$^{+/+}$ LSK cells (C). (D) Representative FACS analysis of engrafted LSK subpopulation 2 months after the transplantation (E) Fold calculated as the proportions of LSK subpopulations 2 months after transplantation divided by the average of those identified a steady state in 2-month-old mice. (F) Absolute number of LSK cells 16 hours (left panel) or 96 hours (right panel) after PBS (open box) or LPS (hatched box). (G) Proportions of LSK subpopulation in *Gata2*$^{+/+}$ (in blue) and *Gata2*$^{R396Q/+}$ (in red) mice 16 hours after injection of PBS (open box) or LPS (hatched box), fold change between LPS and PBS condition is shown at the top of the boxes. (H) Absolute number of the specific subpopulations of LSK from *Gata2*$^{R396Q/+}$ and *Gata2*$^{+/+}$ mice 16 hours (left panel) or 96 hours (right panel)after injection of LPS (open square) or PBS (closed square) injected mice. *Gata2*$^{+/+}$ mice are in blue and *Gata2*$^{R396Q/+}$ mice are in red. LSK: Lineage$^{-}$ Kit$^{+}$ Sca-1$^{+}$ cells; LT-: long-term- and ST-: short-term- HSC: hematopoietic cells; MPP: Multi-Potent Progenitor.. LPS: Lipopolysaccharide. Each dot represents an individual mouse. Results are shown as mean ± SD; Unpaired t-test. *p<0.05, **p<0.01, ***p<0.001, ****p<0.0001, ns: non-significant.

#### Figure S6

(A) Representation of protein-protein interaction network of the DEGs by Metascape analysis. The right panel indicates the proportion *Gata2*$^{+/+}$ (in blue) and *Gata2*$^{R396Q/+}$ (in red). (B) GSEA enrichment of neutrophil mediated immunity pathway in LSK. (C) Transcription factor (TF) activity modulation in *Gata2*$^{R396Q/+}$ compared to *Gata2*$^{+/+}$ LSK using DiffTF algorithm. Enhanced activity is indicated in pink, lower activity in blue. Activating, repressing and undetermined TF were indicated by color code. (D) Pie charts illustrating the distribution of ATAC-seq consensus peaks relative to gene features for each cluster defined in Figure 5G. (E) Expression of *Smad* genes in *Gata2*$^{+/+}$ cells (blue) and *Gata2*$^{R396Q/+}$ (red). (F) Expression of *Erg*, *fli1* and *Gata2* gene in *Gata2*$^{+/+}$ (blue) and *Gata2*$^{R396Q/+}$ LSK cells. For *Gata2*, the wild type allele is indicated with a plain histogram while the mutant allele expression is indicated with hatched histogram. (G) Genome track with exons in black and 3'UTR and 5'UTR in grey of the *Smad1* and *Gata2* gene indicating on the top panel a higher differential accessibility (in red) or lower differential accessibility (in blue) of the different locus. In the middle panel is indicated the position of Gata2, Fli1 and Erg binding sites according to ReMap2022.

#### Figure S7

(A) Gata2 stain index defined as [[Geometric Mean (population of interest) – Geometric Mean (negative population)] / 2* Standard deviation (negative population)] on the different LSK subpopulations on *Gata2*$^{+/+}$ (in bleu) *Gata2*$^{R396Q/+}$ (in red). Differential between SI of mutated cells and SI of unmutated cells is indicated on the graph for each population. (B) Dot plots representing the intracellular FACS analyses of Gata1 proteins on the different LSK subpopulations with CD48 on the x-axis. (C) Gata2 and Gata1 stain index on the different LSK subpopulations on *Gata2*$^{+/+}$ (in bleu) *Gata2*$^{R396Q/+}$ (in red). (D) Decomposed heatmap showing the average fold in LT-HSC, ST-HSC or MPP3-4 of *Gata2*$^{+/+}$ (in bleu) *Gata2*$^{R396Q/+}$ (in red) LSK subpopulations. First panel shows common genes up or downregulated in mutated LT-HSC, ST-HSC and MPP3-4 subpopulations. Second panel shows differential genes only in mutated LT-HSC population (up (top) or down-regulated (bottom)). Third panel shows genes up (top) or down (bottom) regulated preferentially in mutated MPP3/4 cells. Fourth panel shows genes not downregulated in MPP3/4(top) or in ST-HSC (bottom panel). Z-score is indicated as a color code. (E) Expression of *Smad6*, *Aqp9*, *Gpc3* and *Cebpa* in LT-, ST-HSC and MPP3-4 of *Gata2*$^{+/+}$ (in bleu) *Gata2*$^{R396Q/+}$ (in red) mice. (F) Expression of *Cd82*, *Vwf*, *Ly6a, Procr* from the Molo signature and of (G) *Muc13* and *Nkg7* of the NoMo signature of LT-HSC from *Gata2*$^{+/+}$ (in bleu) *Gata2*$^{R396Q/+}$ (in red) mice. (H) Cell type annotation as referred on the left panel. On the right *Igf1*, *Sfrp2* and *S1pr3* expression in BM cells of young mice assessed by scRNA-Seq (Baccin et al. 2020)[66]. For detailed hematopoietic cell type annotation, refer to https://nicheview.shiny.embl.de.

#### Figure S8

(A) TriMap on bone marrow cells of PBS- or LPS- injected *Gata2*$^{+/+}$ (in bleu) and *Gata2*$^{R396Q/+}$ (in red) mice depicting the different HSPC subpopulations showing the population density and the intensity of the CD16/32, CD48, Kit, Flt3, CD61 and CD34 markers on each population. Intensity of each marker is depicted following a colored scale. (B) Density plot showing the CD150, CD41 and Ki67 staining on cells of *Gata2*$^{+/+}$ and *Gata2*$^{R396Q/+}$ LPS-injected mice. (C) Expression of Gata2 in young and old wild-type LT-HSC from Maryanovich et al. data[52] (D) TriMap on the different progenitor subpopulations of middle-aged mice showing the population density and the intensity of the CD48, CD34, Kit, Sca-1 and Flt3 markers on each population following a colored scale. (E) Number of total CFU or CFU-GEMM or -Mk obtained from LT-HSC of *Gata2*$^{+/+}$ (blue) and Gata2$^{R396/+}$ (red) middle-aged mice after clonogenic assay. (F) May-Grunwald Giemsa coloration of cells after methylcellulose cultures of LT-HSC from *Gata2*$^{+/+}$ (left panel) and *Gata2*$^{R396Q/+}$ (right panel) middle-aged mouse. E: Erythroid cells. M: monocyte, Ma: Macrophage, Mk: Megakaryocytes. (G) Genomic analysis of one GATA2 deficient patient diagnosed with AML. Circos plot with structural variations (left panel). Visualization on IGV viewer of the variant allele frequency (VAF) after Whole Genome Sequencing (WGS) of infiltrated bone marrow (first line) and healthy tissue (third line) or Whole Exome Sequencing of infiltrated bone marrow (second line). VAF at the RNA level is visualized after RNA-Seq analysis (last line).

CFU: Colony forming unit; GEMM: Granulocyte, erythroid, macrophage, megakaryocyte; Mk: Megakaryocytes. Each dot represents an individual mouse.

#### Table S1

Listing of *GATA2* germline mutations reported in 11 previous studies[6,8,10,12,16,31–36].

#### Table S2

Chi-square on mice obtained after four breeding between *Gata2*$^{+/-}$ and *Gata2*$^{R396Q/+}$ mice. Are shown expected and observed numbers and percentages of living pups. *p*-value (two-tailed) is 0.0117 with 3 degrees of freedom. Chi-square 11.00.

### Supplemental Figure 1

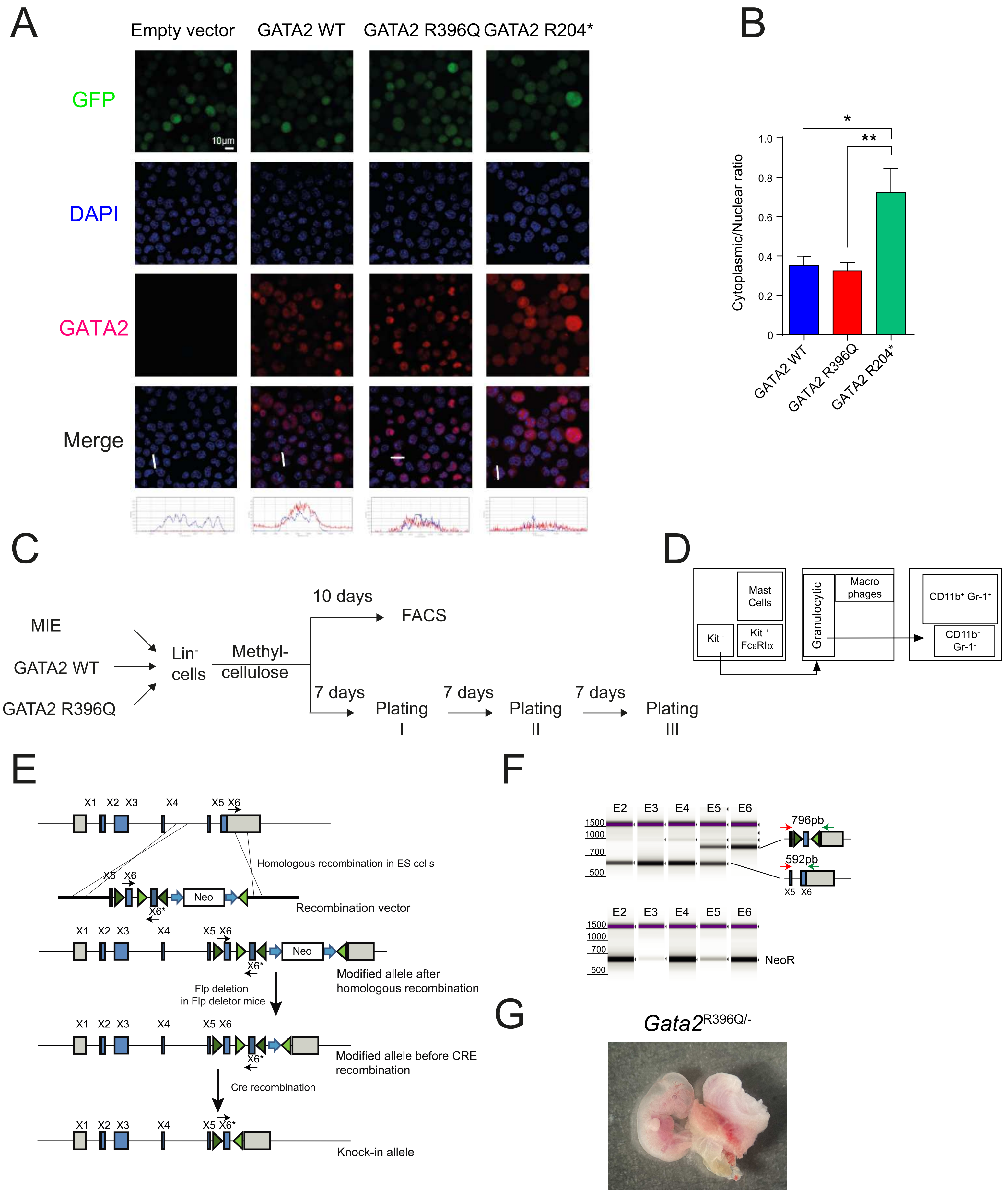

H

E12.5

| Embryo | 1 | 2 | 3 | 4 | 5 | 6 | 7 | 8 | 9 | 10 | 11 | 12 | 13 | 14 | 15 | 16 | 17 | 18 |
|---|---|---|---|---|---|---|---|---|---|---|---|---|---|---|---|---|---|---|
| Wt allele | Y | Y | Y | Y | Y | N | Y | Y | Y | Y | Y | Y | Y | Y | N | Y | Y | N |
| R396Q allele | N | N | N | N | Y | Y | N | N | N | N | N | N | Y | Y | Y | N | N | Y |
| Neo allele | N | Y | N | Y | N | Y | Y | N | N | Y | Y | Y | N | N | Y | N | N | Y |
| Genotype Gata2 | +/+ | +/- | +/+ | +/- | R396Q/+ | R396Q/- | +/- | +/+ | +/+ | +/- | +/- | +/- | R396Q/+ | R396Q/+ | R396Q/- | +/+ | +/+ | R396Q/- |

### Supplemental Figure 2

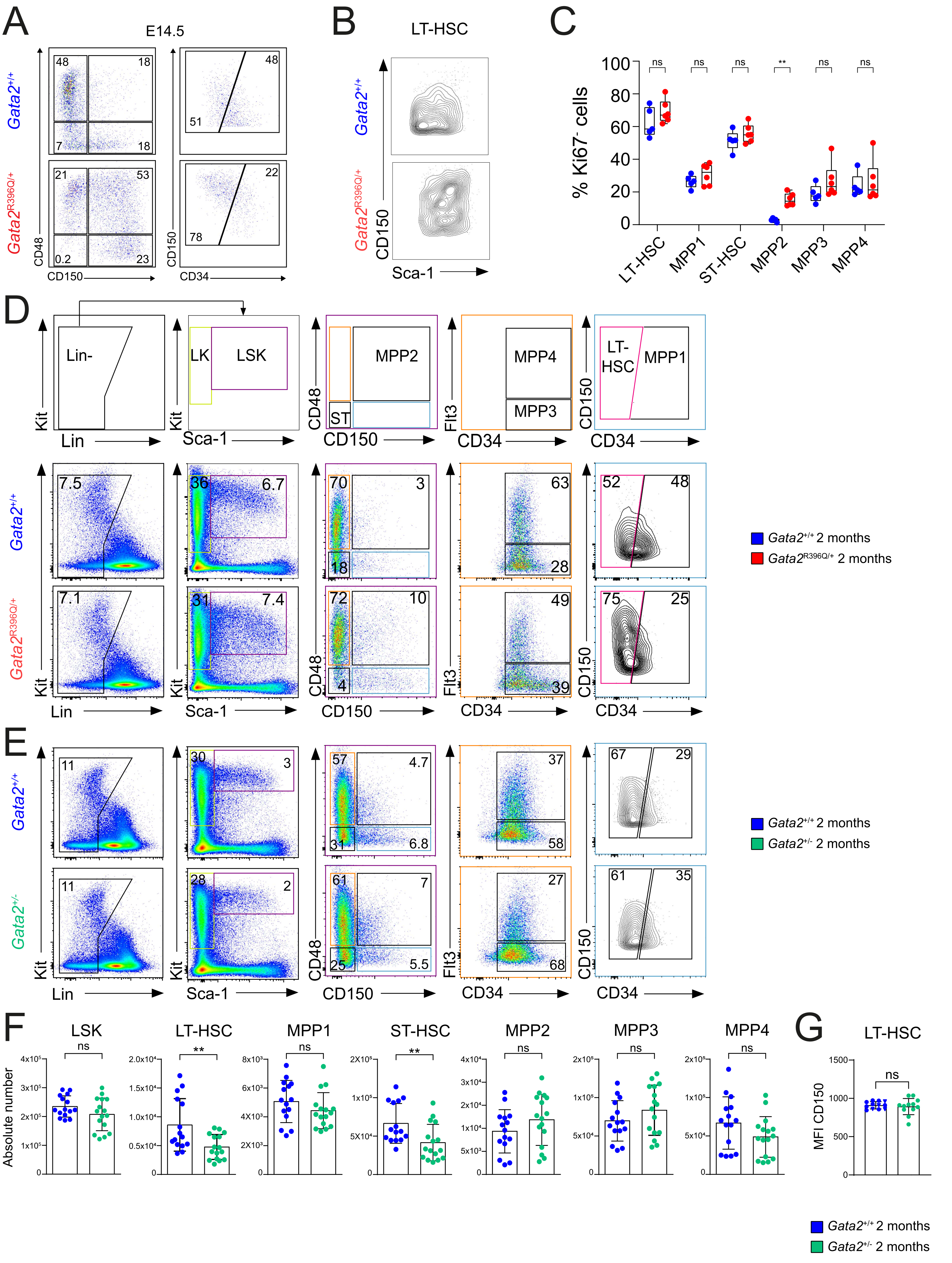

### Supplemental Figure 3

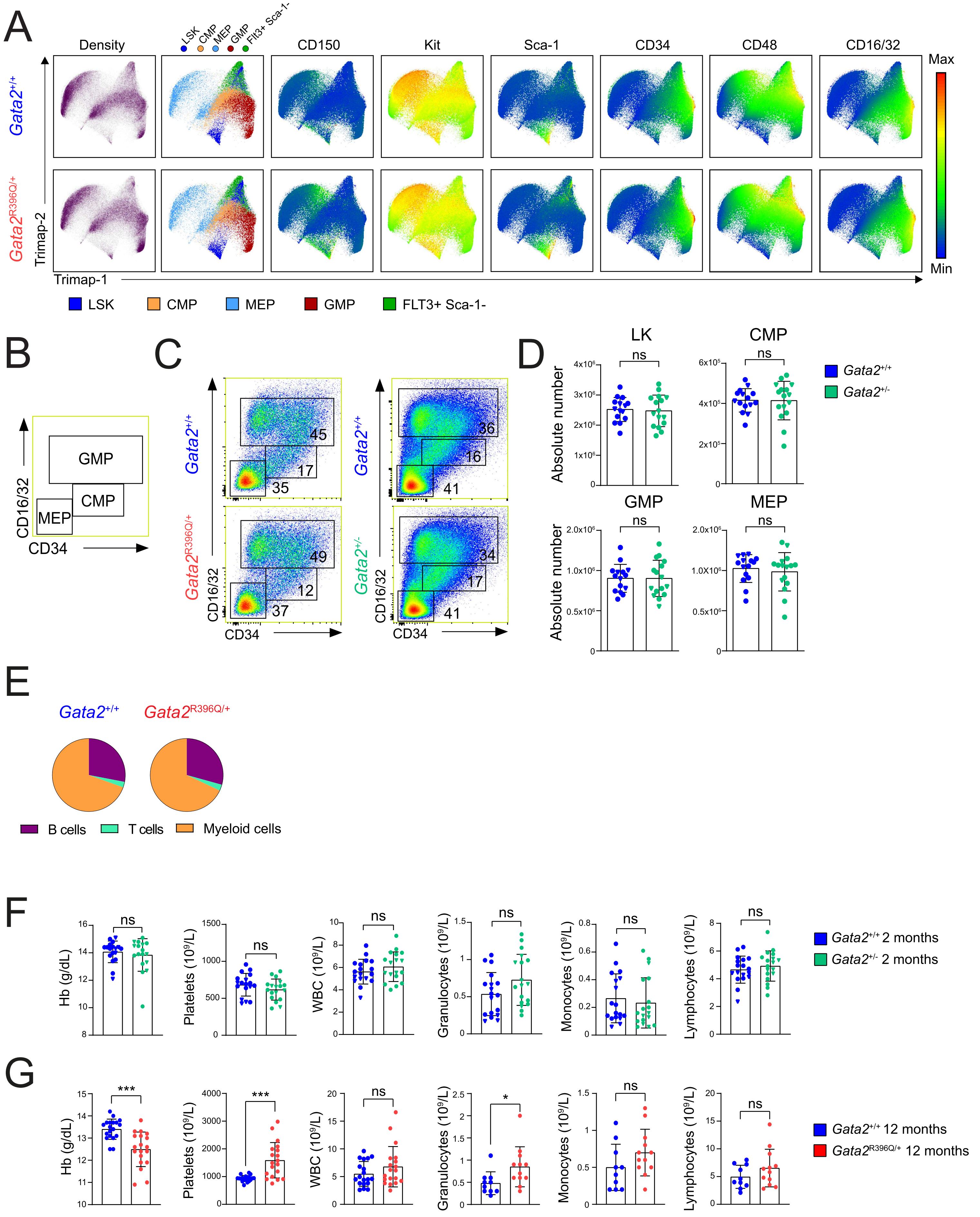

### Supplemental Figure 4

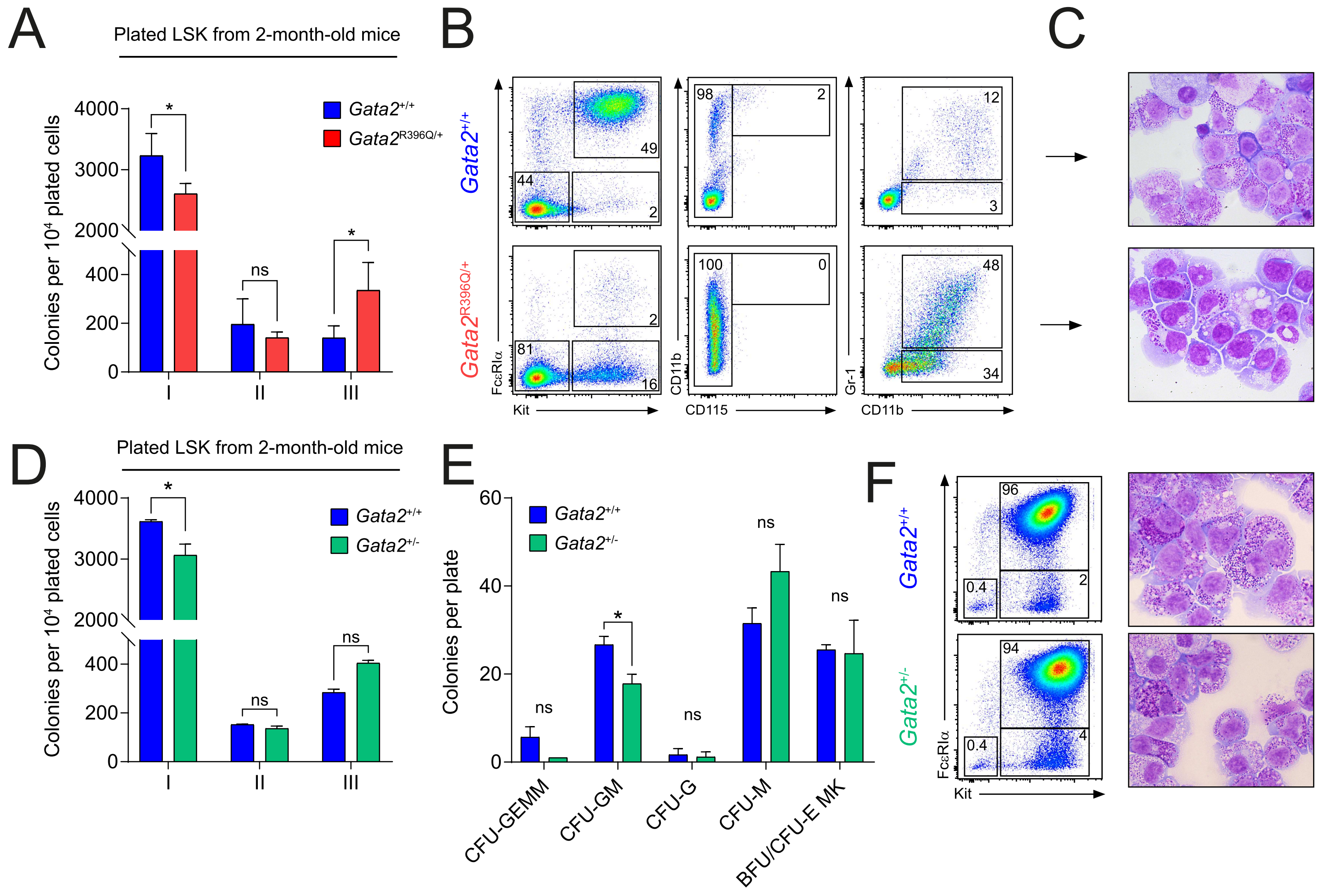

### Supplemental Figure 5

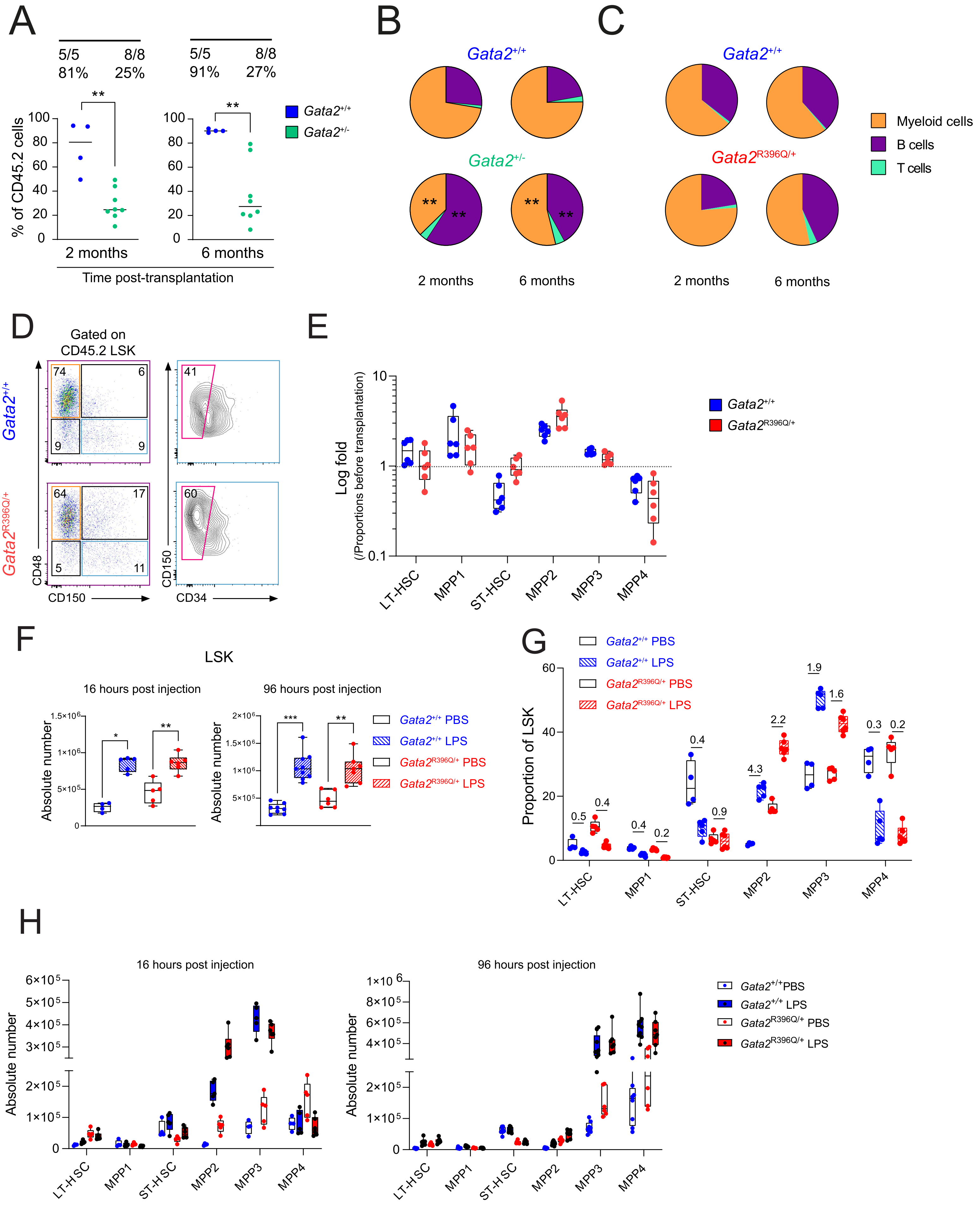

### Supplemental Figure 6

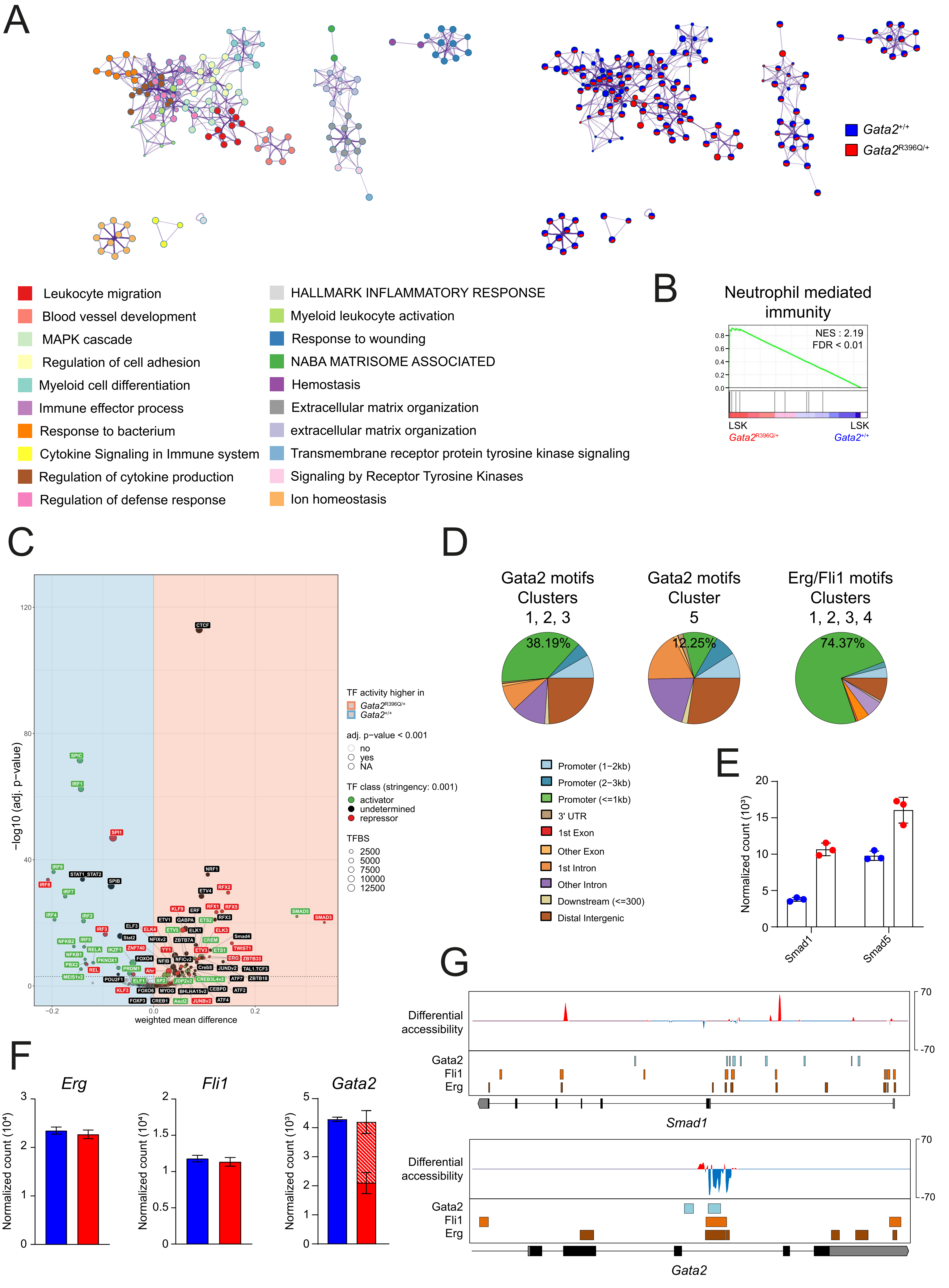

### Supplemental Figure 7

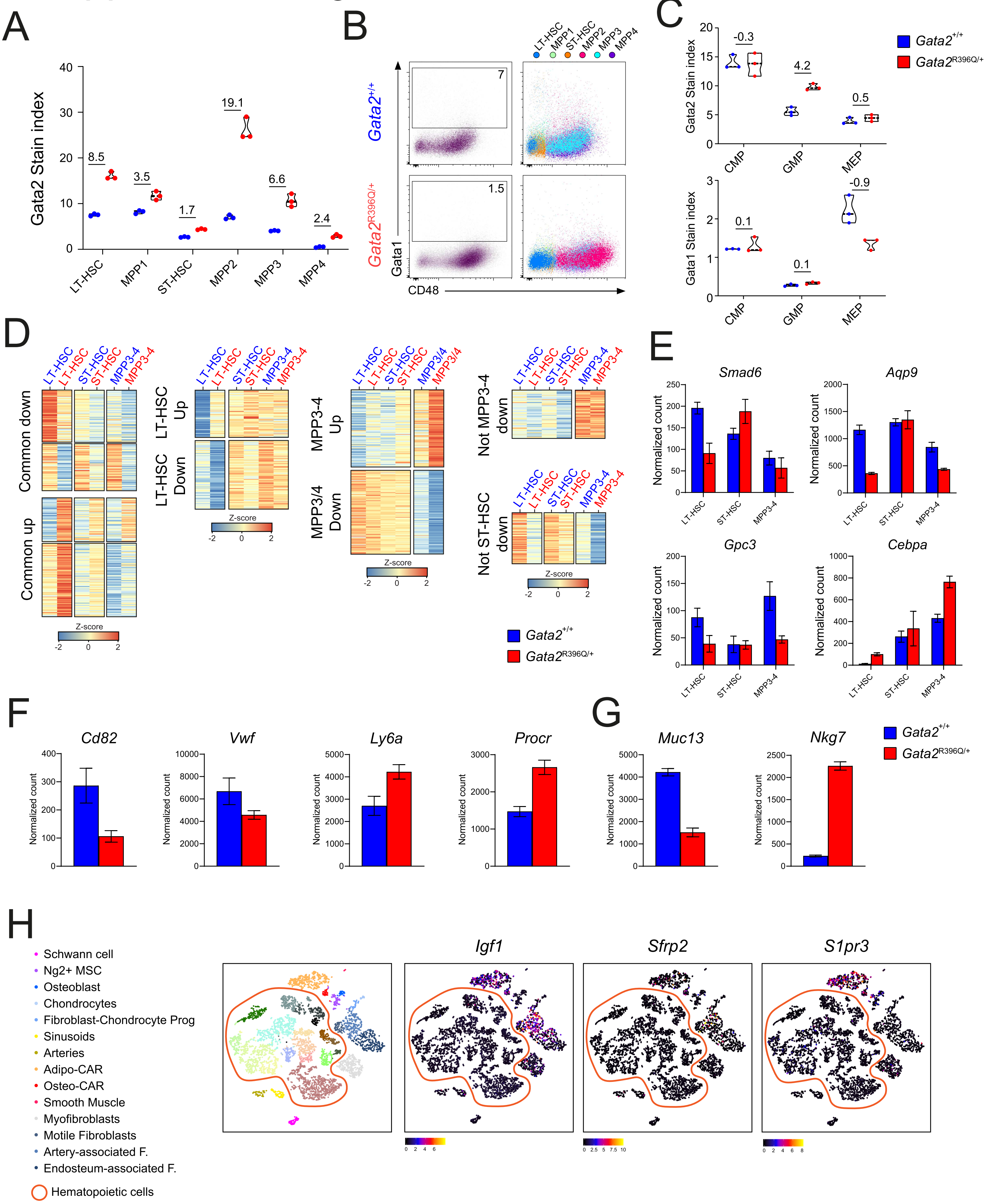

### Supplemental Figure 8

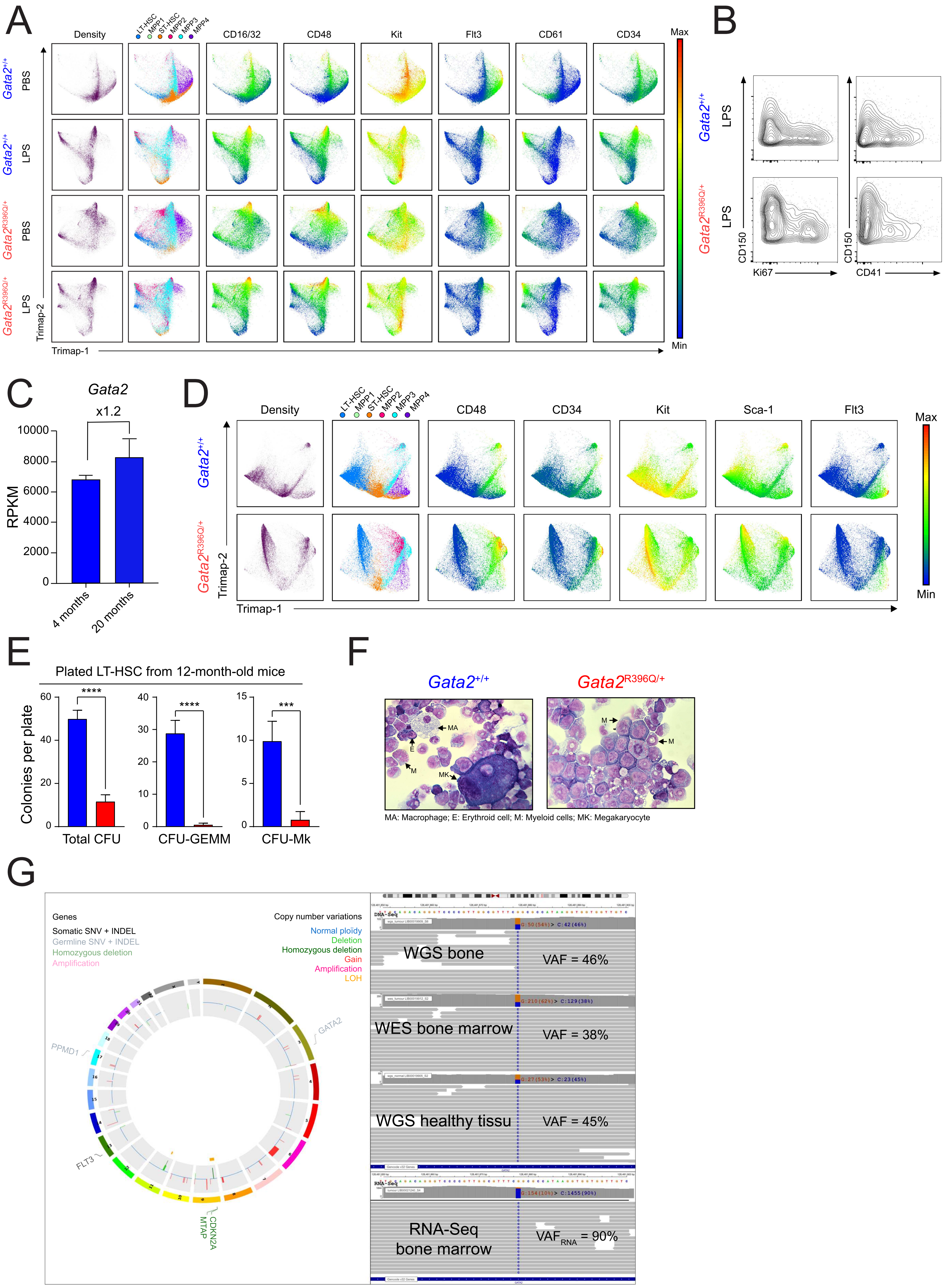

**Table S1.**

| Article/French Registry | Mutation | Codon | Type |
|---|---|---|---|
| Bödor et al | p.Thr354Met | 354 | Missense |
| Burak | p.Arg337* | 337 | Nonsense |
| Hahn et al. | p.Ala341_Gly346del | 341 | Inframe |
| Hahn et al. | p.Thr354Met | 354 | Missense |
| Hahn et al. | p.Thr355del | 355 | Inframe |
| Hahn et al. | p.Leu359Val | 359 | Missense |
| Hahn et al. | p.Arg362Gln | 362 | Missense |
| Holme et al. | p.Leu105Profs*15 | 105 | Frameshift |
| Holme et al. | p.Thr354Met | 354 | Missense |
| Holme et al. | p.Arg396Glu | 396 | Missense |
| Hsu et al. | p.Asp367Glyfs*17 | 367 | Frameshift |
| Hsu et al. | p.Met388Trp | 388 | Missense |
| Hsu et al. | p.Arg396Q | 396 | Missense |
| Hsu et al. | p.Arg398Trp | 398 | Missense |
| Nakazawa | p.Lys378Asnfs*12 | 378 | Frameshift |
| Oleaga-Quintas | p.Ser201* | 201 | Nonsense |
| Oleaga-Quintas | p.Ser201* | 201 | Nonsense |
| Oleaga-Quintas | p.Thr306Alafs*77 | 306 | Frameshift |
| Oleaga-Quintas | p.Arg330* | 330 | Nonsense |
| Oleaga-Quintas | p.Ser340Lysfs*40 | 340 | Frameshift |
| Oleaga-Quintas | p.Ala342Profs*45 | 342 | Frameshift |
| Oleaga-Quintas | p.Gly346Serfs*40 | 346 | Frameshift |
| Oleaga-Quintas | p.Asp367Glyfs*17 | 367 | Frameshift |
| Oleaga-Quintas | p.Met388Thr | 388 | Missense |
| Oleaga-Quintas | p.Met388Thr | 388 | Missense |
| Oleaga-Quintas | p.Arg396Gln | 396 | Missense |
| Oleaga-Quintas | p.Arg396Trp | 396 | Missense |
| Oleaga-Quintas | p.Arg398Trp | 398 | Missense |
| Oleaga-Quintas | p.Arg398Gln | 398 | Missense |
| Oleaga-Quintas | p.Arg398Trp | 398 | Missense |
| Ostergaard et al. | p.Arg78Profs*107 | 78 | Frameshift |
| Ostergaard et al. | p.Leu105Profs*15 | 105 | Frameshift |
| Ostergaard et al. | p.Ala194Serfs*8 | 194 | Frameshift |
| Ostergaard et al. | p.Arg337* | 337 | Nonsense |
| Ostergaard et al. | p.Ala341Argfs*38 | 341 | Frameshift |
| Ostergaard et al. | p.Ala341Profs*45 | 341 | Frameshift |
| Ostergaard et al. | p.Arg361Leu | 361 | Missense |
| Ostergaard et al. | p.Cys373Arg | 373 | Missense |
| French Registry | p.Val47_Gly480delinsGlyLe | 47 | Frameshift |
| French Registry | p.Ser106Cysfs*78 | 106 | Frameshift |
| French Registry | p.Val118Glyfs*100 | 118 | Frameshift |
| French Registry | p.Tyr141* | 141 | Nonsense |
| French Registry | p.Gly146Valfs*72 | 146 | Frameshift |
| French Registry | p.Glu180* | 180 | Nonsense |
| French Registry | p.Ser201* | 201 | Nonsense |
| French Registry | p.Arg204* | 204 | Nonsense |
| French Registry | p.Arg224* | 224 | Nonsense |
| French Registry | p.His243Profs*38 | 243 | Frameshift |
| French Registry | p.Trp306Alafs*77 | 306 | Frameshift |
| French Registry | p.Tyr314Cysfs*66 | 314 | Frameshift |
| French Registry | p.Gly327Glufs*58 | 327 | Frameshift |
| French Registry | p.Arg330* | 330 | Nonsense |
| French Registry | p.Ala342Argfs*42 | 342 | Frameshift |
| French Registry | p.Ala342Profs*45 | 342 | Frameshift |
| French Registry | p.Arg344Glyfs*43 | 344 | Frameshift |
| French Registry | p.Cys349Arg | 349 | Missense |
| French Registry | p.Cys349Tyr | 349 | Missense |
| French Registry | p.Cys349Leufs*35 | 349 | Frameshift |
| French Registry | p.Thr354Pro | 354 | Missense |
| French Registry | p.Thr354Arg | 354 | Missense |
| French Registry | p.Thr354Met | 354 | Missense |
| French Registry | p.Thr354Met | 354 | Missense |
| French Registry | p.Thr355Aspfs*29 | 355 | Frameshift |
| French Registry | p.Thr357I | 357 | Missense |
| French Registry | p.Thr358Asn | 358 | Missense |
| French Registry | p.Leu359Ser | 359 | Missense |
| French Registry | p.Trp360-Arg361dup | 360 | Inframe |
| French Registry | p.Trp360* | 360 | Nonsense |
| French Registry | p.Arg361Gly | 361 | Missense |
| French Registry | p.Arg361Cys | 361 | Missense |
| French Registry | p.Arg361His | 361 | Missense |
| French Registry | p.Arg361His | 361 | Missense |
| French Registry | p.Arg361His | 361 | Missense |
| French Registry | p.Arg362* | 362 | Nonsense |
| French Registry | p.Arg362* | 362 | Nonsense |
| French Registry | p.Arg362* | 362 | Nonsense |
| French Registry | p.Arg362* | 362 | Nonsense |
| French Registry | p.Arg362Pro | 362 | Missense |
| French Registry | p.Asp367Glyfs*17 | 367 | Frameshift |
| French Registry | p.Pro368Argfs*15 | 368 | Frameshift |
| French Registry | p.Ala372Val | 372 | Missense |
| French Registry | p.Ala372Thr | 372 | Missense |
| French Registry | p.Ala372Thr | 372 | Missense |
| French Registry | p.Ala372Thr | 372 | Missense |
| French Registry | p.Cys373Tyr | 373 | Missense |
| French Registry | p.Leu375Pro | 375 | Missense |
| French Registry | p.Asn381Metfs*6 | 381 | Frameshift |
| French Registry | p.Pro385Gln | 385 | Missense |
| French Registry | p.Met388Val | 388 | Missense |
| French Registry | p.Arg396Trp | 396 | Missense |
| French Registry | p.Arg396Trp | 396 | Missense |
| French Registry | p.Arg396Gln | 396 | Missense |
| French Registry | p.Arg398Trp | 398 | Missense |
| French Registry | p.Arg398Gln | 398 | Missense |
| French Registry | p.Arg398Gln | 398 | Missense |
| French Registry | p.Arg398Gln | 398 | Missense |
| French Registry | p.Arg398Gln | 398 | Missense |

| Article/French Registry | Mutation | Codon | Type |
|---|---|---|---|
| Spinner et al. | p.Gln83Profs*102 | 83 | Frameshift |
| Spinner et al. | p.Gly101Alafs*16 | 101 | Frameshift |
| Spinner et al. | p.Val140Cysfs*44 | 140 | Frameshift |
| Spinner et al. | p.Gly199Leufs*21 | 199 | Frameshift |
| Spinner et al. | p.Tyr260Cysfs*25 | 260 | Frameshift |
| Spinner et al. | p.Ala318Thrfs*12 | 318 | Frameshift |
| Spinner et al. | p.Arg330* | 330 | Nonsense |
| Spinner et al. | p.Leu332Thrfs*53 | 332 | Frameshift |
| Spinner et al. | p.Arg337* | 337 | Nonsense |
| Spinner et al. | p.Thr354Met | 354 | Missense |
| Spinner et al. | p.Thr354Met | 354 | Missense |
| Spinner et al. | p.Thr354Met | 354 | Missense |
| Spinner et al. | p.Thr354Met | 354 | Missense |
| Spinner et al. | p.Arg361Cys | 361 | Missense |
| Spinner et al. | p.Arg361del | 361 | Inframe |
| Spinner et al. | p.Asp367Glyfs*15 | 367 | Frameshift |
| Spinner et al. | p.Asn371Lys | 371 | Missense |
| Spinner et al. | p.Cys373_Tyr377del | 373 | Inframe |
| Spinner et al. | p.Met388Thr | 388 | Missense |
| Spinner et al. | p.Arg396Trp | 396 | Missense |
| Spinner et al. | p.Arg396Trp | 396 | Missense |
| Spinner et al. | p.Arg396Gln | 396 | Missense |
| Spinner et al. | p.Arg396Gln | 396 | Missense |
| Spinner et al. | p.Arg396Gln | 396 | Missense |
| Spinner et al. | p.Arg396Gln | 396 | Missense |
| Spinner et al. | p.Arg398Trp | 398 | Missense |
| Spinner et al. | p.Arg398Trp | 398 | Missense |
| Spinner et al. | p.Arg398Trp | 398 | Missense |
| Spinner et al. | p.Arg398Trp | 398 | Missense |
| Spinner et al. | p.Arg398Trp | 398 | Missense |
| Wang et al. | p.Gly268* | 268 | Nonsense |
| Wang et al. | p.Cys298Leufs*86 | 298 | Frameshift |
| Wlodarski et al. | p.Ser54* | 54 | Nonsense |
| Wlodarski et al. | p.Val70Leufs*114 | 70 | Frameshift |
| Wlodarski et al. | p.Ala103fs*116 | 103 | Frameshift |
| Wlodarski et al. | p.139Cysfs*45 | 139 | Frameshift |
| Wlodarski et al. | p.Gly200Valfs*18 | 200 | Frameshift |
| Wlodarski et al. | p.Gly200Valfs*18 | 200 | Frameshift |
| Wlodarski et al. | p.Ser201* | 201 | Nonsense |
| Wlodarski et al. | p.Val211Argfs*72 | 211 | Frameshift |
| Wlodarski et al. | p.Leu229Cysfs*5 | 229 | Frameshift |
| Wlodarski et al. | p.Gly268* | 268 | Nonsense |
| Wlodarski et al. | p.His323Glnfs*61 | 323 | Frameshift |
| Wlodarski et al. | p.Leu332Glnfs*60 | 332 | Frameshift |
| Wlodarski et al. | p.Arg337* | 337 | Nonsense |
| Wlodarski et al. | p.Arg344Lysfs*37 | 344 | Frameshift |
| Wlodarski et al. | p.Thr347Argfs*38 | 347 | Frameshift |
| Wlodarski et al. | p.Cys348Phe | 348 | Missense |
| Wlodarski et al. | p.Cys352Gly | 352 | Missense |
| Wlodarski et al. | p.Thr356_Asn365del | 356 | Inframe |
| Wlodarski et al. | p.Thr357Ala | 357 | Missense |
| Wlodarski et al. | p.Arg361Cys | 361 | Missense |
| Wlodarski et al. | p.Arg361His | 361 | Missense |
| Wlodarski et al. | p.Arg362* | 362 | Nonsense |
| Wlodarski et al. | p.Arg362* | 362 | Nonsense |
| Wlodarski et al. | p.Asn371Lys | 371 | Missense |
| Wlodarski et al. | p.Leu375Profs*12 | 375 | Frameshift |
| Wlodarski et al. | p.Lys390Glu | 390 | Missense |
| Wlodarski et al. | p.Arg396Gln | 396 | Missense |
| Wlodarski et al. | p.Arg396Gln | 396 | Missense |
| Wlodarski et al. | p.Arg396Trp | 396 | Missense |
| Wlodarski et al. | p.Arg396Trp | 396 | Missense |
| Wlodarski et al. | p.Arg396Trp | 396 | Missense |
| Wlodarski et al. | p.Arg398Trp | 398 | Missense |
| Wlodarski et al. | p.Gly415Lys | 415 | Missense |

**Table S2.**

| Phenotypes | Observed # | Expected # | Expected % | Observed % |
|---|---|---|---|---|
| $Gata2^{+/+}$ | 8 | 6 | 25,00 | 33,33 |
| $Gata2^{+/-}$ | 11 | 6 | 25,00 | 45,83 |
| $Gata2^{R396Q/+}$ | 5 | 6 | 25,00 | 20,83 |
| $Gata2^{R396Q/-}$ | 0 | 6 | 25,00 | 0,0 |
| TOTAL | 24 | 24 | 100,0 | 100,00 |

Chi-square test

Chi-square 11.00

DF 3

P value (two-tailed) 0.0117

P value summary *

Is discrepancy significant (P < 0.05)? Yes
